# Distinct and overlapping correlates of fear acquisition and extinction across different neuroimaging modalities

**DOI:** 10.64898/2026.08.24.746689

**Authors:** Arslan Gabdulkhakov, Christoph Fraenz, Dorothea Metzen, Julian Packheiser, Christian J. Merz, Nikolai Axmacher, Erhan Genç

## Abstract

Interindividual differences in fear acquisition and extinction have been related to variation in specific brain correlates. However, variability in experimental setups complicates the integration of findings. Here, we present a combined fear acquisition (n = 101) and extinction (n = 88) study from which we obtained distinct microstructural, macrostructural, and connectivity brain properties with magnetic resonance imaging in healthy, young participants. The properties included regional brain volume, cortical surface area and thickness, neurite density and orientation dispersion, and nodal efficiency of structural and functional connectivity. Fear responses were quantified as changes in skin conductance. Data from 360 cortical and 16 subcortical brain regions as well as the efficiency of network connections between them were used as independent variables in bootstrapped and cross-validated regularized regression models. Results show that numerous brain regions, spanning the so-called ‘fear and extinction network’ and extending beyond it, contribute to fear acquisition, to extinction, or dynamically shift between both phases. For several brain regions, data from multiple imaging modalities showed a high degree of concordance for several brain regions. These findings call for further research to examine the potential interplay between brain correlates shaping fear acquisition and extinction, as opposed to studying the imaging modalities in isolation.

**Highlights:**

- Multimodal brain properties relate to individual differences in fear learning
- Fear acquisition showed stronger cross-modal convergence than extinction
- Hippocampal, prefrontal, and sensory regions related to fear acquisition
- Extinction involved distributed frontoparietal and cingulo-opercular markers
- Acquisition and extinction shared few stable neural correlates

## 1. Introduction

Fear is a fundamental emotion that plays a crucial role in human survival and adaptation. The acquisition and extinction of fear are essential processes through which individuals learn to respond to and regulate threatening stimuli in their environment (Craske et al., 2006). Fear acquisition refers to the initial learning of fear associations, wherein a typically neutral stimulus is associated with an aversive event and becomes a conditioned stimulus (CS), leading to the subsequent elicitation of fear responses. On the other hand, fear extinction involves the gradual reduction and eventual cessation of fear responses towards the CS when the previously learned fear association is no longer relevant or threatening (Lonsdorf et al., 2017). Individuals show significant differences in their susceptibility to acquiring and extinguishing fear responses (Lonsdorf & Merz, 2017). Recently, elucidating the neurobiological underpinnings of these inter-individual differences has become a major focus in neuroscience. While previous research has identified various brain regions where specific neuronal parameters correlate with an individual’s responsiveness to fear (e.g., MacNamara et al., 2015; Ehlers et al., 2020), these potential correlates are typically analyzed in isolation. However, a recent large-sample study by Gomes et al. (2026) demonstrated the value of a multimodal approach, showing that functional, effective, and structural connectivity jointly predict individual differences in fear acquisition and extinction within a predefined set of regions of interest. Still, it remains an open question whether the neurobiology of fear acquisition and extinction is characterized by a multimodal set of brain properties across the whole brain, and whether microstructural tissue properties provide additional predictive information.

Investigating the neural correlates of fear acquisition and extinction relies on controlled experiments that typically employ classical conditioning paradigms (Lonsdorf et al., 2017). In typical experiments in humans, a stimulus (such as a tone or a visual cue) is contingently paired with an aversive unconditioned stimulus (US; such as a mild electric shock or an aversive image), causing the stimulus to become a conditioned stimulus (CS+) that elicits a conditioned fear response. During fear acquisition training, another stimulus (CS–) is never paired with the US, and thereby marks its absence (Lonsdorf et al., 2017). Conditioned fear responses (CR) can be quantified via physiological measures (such as skin conductance responses, SCRs) and self-report. Extinction training aims to diminish previously established CR by presenting the CS+ without the aversive event, leading to a gradual reduction in fear.

Numerous studies have explored the neural correlates of fear acquisition and extinction using functional magnetic resonance imaging (fMRI). These investigations have provided valuable insights into the underlying brain regions associated with these processes. For example, the amygdala, a key region involved in emotional processing, has been shown to exhibit heightened activation during fear acquisition, primarily during the early phase (Buchel et al., 1999; LaBar et al., 1998; Phelps, 2004). The amygdala’s engagement in fear acquisition has been debated in more recent studies (Visser et al., 2021; Wen et al., 2022, 2024) but is supported by its connectivity with other regions such as the prefrontal cortex (PFC), insula, and hippocampus, which play critical roles in the evaluation, regulation, and contextualization of fear responses (Milad & Quirk, 2012; but see Fullana et al., 2019). Milad, Wright, et al. (2007) demonstrated that the ventromedial PFC (vmPFC) is deactivated during fear acquisition, activated during fear extinction and that the vmPFC activation during extinction recall positively correlated with extinction retention. Recent meta-analyses have identified whole brain networks showing increased activation in response to fear- or safety-related stimuli (Fullana et al., 2016, 2018).

While much research has focused on task-based brain activation during fear acquisition and extinction, recent studies have started exploring the relationship between resting-state brain connectivity and fear conditioning. For example, it has been demonstrated that fear acquisition and extinction can alter the resting-state connectivity between various brain regions (Feng et al., 2013, 2014; Fraenz et al., 2025; Hermans et al., 2016; Martynova et al., 2020; Schultz et al., 2012). Respective connections were, among others, constituted by the amygdala, dorsal anterior cingulate cortex (dACC), vmPFC, insula, and hippocampus. Resting-state brain connectivity is likely to change as a result of fear conditioning, but it can also, at least to some extent, predict the effect of fear conditioning. For example, resting-state connectivity between amygdala and vmPFC, measured after fear acquisition, is positively correlated with the effect of subsequent fear extinction (Feng et al., 2016). Beyond pairwise connectivity measures, graph-theoretical metrics such as the nodal efficiency, which captures the communicative centrality of individual regions, have been used to predict interindividual differences in cognitive and emotional processing from resting-state fMRI data (Genç et al., 2019, 2023; Metzen et al., 2024). These measures thus represent a promising framework for characterizing the network-level architecture that may underlie individual differences in fear acquisition and extinction.

In addition to task-based and resting-state fMRI studies, previous studies have explored the structural correlates of fear acquisition and extinction using structural MRI techniques. For instance, cortical thickness of the posterior insula (Hartley et al., 2011) and the dorsal anterior cingulate cortex (Milad, Quirk, et al., 2007) has been shown to be positively associated with fear responses evoked in a classical fear conditioning paradigm. Following a similar approach, a recent study investigating fear learning in anxiety patients and healthy controls found cortical thickness of the dorsomedial and dorsolateral PFC to be negatively associated with SCRs (Abend et al., 2020). In view of subcortical regions, it has been demonstrated that left amygdala volume is positively correlated with the magnitude of conditioned fear responses, whereas bilateral hippocampal volumes are associated with the magnitude of differential contingency ratings, i.e., how well participants can remember which CS was paired with the US and which was not (Cacciaglia et al., 2015). Furthermore, larger hippocampal volume was shown to be indicative of stronger SCRs to CS+ presentations during fear acquisition (Pohlack et al., 2012). Right amygdala volume was found to be positively correlated with SCR differences between CS+ and CS− presentations during fear acquisition (Winkelmann et al., 2016).

In addition to studying brain macrostructure and functional brain connectivity, previous research has employed diffusion-weighted imaging (DWI) and fiber tracking techniques to investigate the role of structural brain connectivity in fear learning processes. DWI allows the examination of white matter tracts in the brain, providing insights into the integrity and organization of neural connections (Basser & Pierpaoli, 1996; Le Bihan et al., 2001). For instance, fractional anisotropy (FA), a diffusion-derived index sensitive to white matter microstructural properties, has been associated with the expression of fear extinction and fear renewal (Nees et al., 2019; Hermann et al., 2017). Specifically, higher FA in the hippocampal portion of the cingulum was associated with higher skin conductance responses (SCRs) during the extinction of contextual conditioned fear (Nees et al., 2019) and with stronger fear renewal in the acquisition context (Hermann et al., 2017). Furthermore, it has been demonstrated that patients suffering from post-traumatic stress disorder and trauma-exposed controls show significant differences with regard to white matter integrity of specific brain regions (Kunimatsu et al., 2020). In another study comparing trauma-exposed individuals and trauma-unexposed individuals, DWI was used to extract nodal efficiency, connectivity, and network features of fear-circuitry related brain regions, including the amygdala, orbitofrontal cortex, and vmPFC, hippocampus, insula, and thalamus. By analyzing the DWI-derived measures via a machine learning approach, the authors were successful in distinguishing between the two groups at each stage of recovery (Im et al., 2017).

As our understanding of the neural correlates of fear acquisition and extinction continues to evolve, researchers have been exploring advanced MRI techniques to investigate the microstructural properties of the brain. Alongside the previously discussed methods, such as task-based and resting-state fMRI, regional brain volume, cortical thickness, and structural connectivity measures derived from DWI, there is another promising diffusion MRI technique called Neurite Orientation Dispersion and Density Imaging (NODDI) (Zhang et al., 2012). NODDI provides valuable insights into microstructural properties by characterizing the density of neurites (axons and dendrites) and the dispersion of their orientations. While NODDI is a relatively novel technique that, to the best of our knowledge, has not been applied to investigate human fear acquisition and extinction. Its potential for quantifying microstructural alterations in the brain has been utilized in other fields, e.g., to investigate interindividual differences in cognitive ability (Genç et al., 2018), language processing (Ocklenburg et al., 2018), and hemispheric asymmetries (Schmitz et al., 2019).

Integrating findings from multiple neuroimaging techniques, including functional and structural measures as well as connectivity analyses, allows for a comprehensive examination of the neural correlates of fear acquisition and extinction. Such multi-modal approaches hold promise in unraveling the complex interactions and mechanisms underlying fear-related behaviors. However, it is challenging to integrate results from previous studies, given their diverse approaches. Typically, most of these studies are focused on only one isolated neuroimaging technique (either structural, functional, or diffusion MRI). In most cases, they investigate merely one specific phase of fear conditioning (either fear acquisition or extinction). Moreover, many studies analyze only a limited set of brain regions, typically selected on the basis of their prior involvement in fear learning. Consequently, such region-of-interest approaches may miss effects outside this a priori defined set. In contrast, Gomes et al. (2026) used a multimodal neuroimaging approach to examine fear learning across a broad set of brain regions and connectivity measures, highlighting the value of moving beyond isolated, predefined regions. Finally, different studies may use different methods (SCRs, pupil dilation, heart rate, self-reports) and statistics (reaction towards CS+ or CS− alone, difference between CS+ and CS− reactions) to quantify fear responses.

In the present work, we investigated the neural correlates of individual differences in fear acquisition and extinction across seven neuroimaging modalities within a single, unified study. Specifically, we asked: (i) which brain properties, across macrostructural, microstructural, and connectivity measures, reliably predict the magnitude of conditioned fear responses and their extinction; (ii) whether predictive brain regions are modality-specific or whether certain regions emerge consistently across multiple imaging modalities; and (iii) whether acquisition and extinction engage overlapping or anatomically distinct sets of predictors. To address these questions, we combined functional and structural connectivity analyses with macrostructural markers (cortical thickness, cortical surface area, regional brain volume) and, for the first time in this context, NODDI-derived microstructural indices (neurite density and orientation dispersion). Analyses were conducted in a comparatively large sample (∼100 participants) using a whole-brain, data-driven approach based on the Human Connectome Project multimodal parcellation, thereby avoiding the region-of-interest constraints that have limited previous work (Gomes et al., 2026).

## 2. Methods

### 2.1 Participants

The sample recruited for the current study consisted of 138 young and healthy participants, most of them college students. Several participants had to be excluded from the initially recruited sample due to various reasons: Two participants were removed because of incomplete or corrupted MRI data files. Thirteen participants reported a CS−/US contingency that was equal to or larger than their reported CS+/US contingency, indicating absent contingency awareness, which is a necessary prerequisite to observe conditioned SCRs (Tabbert et al., 2011). Finally, 22 participants had to be removed from the analysis of fear acquisition data, and 35 participants had to be removed from the analysis of fear extinction data because of technical difficulties with the electrical stimulation, corrupted or missing SCR data files, or excessive noise in the participants’ SCR recordings. Hence, all analyses related to fear acquisition were carried out with data from 101 participants (56 women; 18 to 26 years, *M* = 21.97, *SD* = 2.07), while analyses related to fear extinction included data from 88 participants (47 women; 18 to 26 years, *M* = 22.08, *SD* = 2.11). We did not observe significant age differences between male and female participants, neither in the fear acquisition (*t*(99) = 1.696, *p* = .0931) nor in the fear extinction group (*t*(86) = 1.505, *p* = .136).

To control for confounding effects caused by handedness, only right-handed individuals, as measured by the Edinburgh Handedness Inventory (Oldfield, 1971), were recruited. All participants had normal or corrected-to-normal vision and were able to understand the provided written and oral instructions. They were either paid for their participation or received course credit. All participants were naive to the purpose of the study and had no prior experience with the fear conditioning paradigm used for the experiment. Participants reported no history of psychiatric or neurological disorders and matched the standard inclusion criteria for fMRI examinations. The study was approved by the local ethics committee of the Faculty of Psychology at Ruhr University Bochum (application number 327). All participants provided written informed consent prior to participation and were treated in accordance with the Declaration of Helsinki.

Although these sample sizes are comparatively large for a neuroimaging study, the number of participants (observations) substantially exceeded the number of predictors in our regression analyses. To address this high-dimensional setting, we used Elastic Net regularization, which combines the L1 penalty of LASSO with the L2 penalty of ridge regression (Zou & Hastie, 2005). This approach is particularly useful when predictors are numerous and correlated, as it can perform variable selection while retaining groups of correlated predictors. Compared with LASSO, Elastic Net may therefore provide more stable estimates in the presence of multicollinearity (Vabalas et al., 2019).

### 2.2 Fear Acquisition and Extinction Paradigm

On day 1, the participants underwent the structural and diffusion MRI, described in detail in Sections 2.4-2.6. On day 2, all participants completed differential fear acquisition followed by fear extinction training while lying in the fMRI scanner. The stimuli and procedures for these paradigms were modified from Milad, Wright, et al. (2007). Participants were instructed to pay close attention to the images presented and told that an electrical stimulation may or may not be presented during the experiment. Electrical stimulation (1 ms pulses with 50 Hz for a duration of 100 ms) was applied as the US via a constant voltage stimulator (STM2000 BIOPAC systems, CA, USA) along with two electrodes attached to the fingertips of the first and second fingers of the right hand. The intensity of electrical stimulation was adjusted for each participant individually prior to the resting-state scan preceding fear acquisition and extinction training on day 2. For this purpose, electrical stimulation was administered at 30 V and raised in increments of 5 V until participants rated the sensation as very unpleasant but not painful.

Stimuli used for fear acquisition and extinction training were presented using the Presentation software package (Neurobehavioral Systems, Albany, CA) and MR-compatible LCD-goggles (Visuastim Digital, Resonance Technology Inc, Northridge, CA). A picture of an office room with a switched-off desk lamp served as the context image. The CS presentation was implemented by the desk lamp, lighting up in either blue or yellow. At the beginning of each trial, a white fixation cross was presented on a black background for 6.8 - 9.5 seconds. Next, the context image was presented for 1 second, which was immediately followed by the presentation of CS+ or CS− for another 6 seconds. During fear acquisition training, the CS+ was paired with electrical stimulation in 62.5 % of trials (10 out of 16). The stimulation was administered 5.9 seconds after CS+ onset and co-terminated with CS+ offset. Exemplary trials for fear acquisition and extinction training are depicted in Figure 1. Fear acquisition training included a total of 32 trials with CS+ and CS− being presented 16 times each in pseudo-randomized order, while fear extinction training included 16 trials with 8 CS+ and 8 CS− presentations, respectively. The first two trials always consisted of one CS+ and one CS− presentation, and so did the last two trials. During fear acquisition training, the first and last CS+ presentations were always paired with electrical stimulation. There were no trials that presented the same type of CS more than twice in consecutive order. CS+ and CS− presentations were distributed equally across both halves. SCRs were assessed using two Ag/AgCl electrodes filled with isotonic (0.05 NaCl) electrolyte medium placed on the hypothenar eminence right below the fifth finger of the left hand. Data were recorded with a sampling rate of 5,000 Hz using Brain Vision Recorder software (Brain Products GmbH, Munich, Germany).

**Figure 1.**
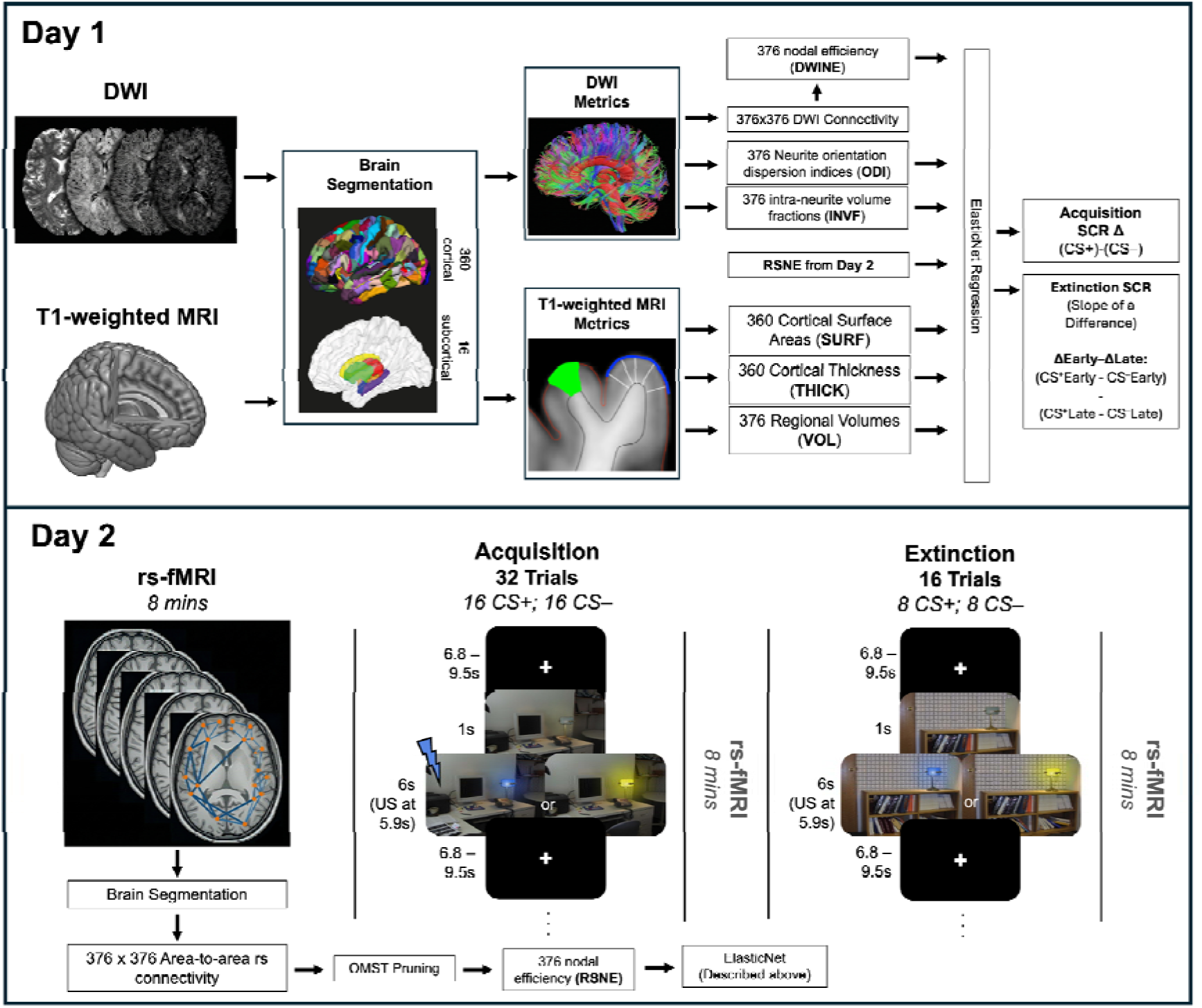
Schematic depiction of data acquisition, preprocessing, and analysis. (Top) Day 1: participants underwent T1-weighted and diffusion-weighted imaging. (Bottom) Day 2: participants underwent a resting-state fMRI (rs-fMRI) scan (used for the extraction of cortical and subcortical connectivity metrics) that was followed by fear acquisition training, another 8-minute rs-fMRI, and extinction training. The two rs-fMRI post-acquisition and extinction training are not analyzed in the present study. At the beginning of each trial, a white fixation cross was presented on a black background for 6.8 - 9.5 seconds. Next, the context image was presented for 1 second, which was followed by the presentation of the CS+ or the CS− for another 6 seconds. During fear acquisition training, the CS+ was paired with electrical stimulation as the US (indicated by a blue lightning bolt) in 62.5 % of trials administered 5.9 seconds after CS+ onset. CS+ and CS− were presented 16 times each in pseudo-randomized order. During fear extinction training, CS+ and CS− were presented 8 times each, but without a US. All brain images obtained prior to fear acquisition and extinction training were parcellated into 360 cortical and 16 subcortical regions. T1-weighted images were used to compute the regional brain volume, cortical surface area, and cortical thickness of each region. Diffusion-weighted images were used to compute measures of neurite density and orientation dispersion for each region. Moreover, fiber tracking was performed to estimate structural connectivity between all possible combinations of brain regions, which were used to compute nodal efficiency. Likewise, fMRI resting-state scans were used to compute average BOLD signal time courses for each region. The resulting metrics were then correlated with each other to obtain estimates of functional connectivity between all possible combinations of brain regions, then pruning of spurious connections, and similarly, the nodal efficiency. Fear acquisition was quantified by subtracting the average SCR to CS− trials from the average SCR to CS+ trials. To quantify extinction learning, CS+ and CS− responses were averaged across the first four (early) and the last four (late) trials, respectively. The difference between average CS+ and CS− responses from the late half was subtracted from the difference between average CS+ and CS− responses from the early half. For each type of brain imaging metric, two regularized regression models were computed. One with the SCR-based measure of fear acquisition as its dependent variable and one with the SCR-based measure of fear extinction.

After fear acquisition and extinction training, participants had to rate the contingencies between CS+, CS−, and US. First, they were asked to estimate the number of electrical stimulations they had received during the respective phase. If they reported having received at least one electrical stimulation, which was to be expected after fear acquisition training, participants were also asked to rate the unpleasantness of the last electrical stimulation on a 9-point Likert scale (1 - “not unpleasant”, 9 - “very unpleasant”) and to estimate at what rates the blue and yellow lamplights had been followed by an electrical stimulation. If participants reported having received no electrical stimulation at all, which was to be expected after fear extinction training, they were merely asked if they had noticed any differences between the two images and how often the blue and the yellow lamplights were presented.

### 2.3 Acquisition and Preprocessing of Skin Conductance Responses

The preprocessing of SCR data was carried out via the Psychophysiological Modelling toolbox in MATLAB (PsPM, version 6.0.0). First, raw data were trimmed and filtered using a median filter with a varying number of time points, such that an optimal number could be found for each participant. Subsequently, artifacts in the trimmed and filtered data were detected using a script based on the automated quality assessment procedure developed by Kleckner et al. (2017). Next, each dataset was manually checked for residual artifacts that may have been missed by the automated artifact detection tool. If any were found, PsPM was used to manually mark all remaining artifacts for subsequent removal. This step was carried out by three researchers with extensive experience in preprocessing SCR data. Participants with an excessive number of artifacts in their data were excluded from further statistical analysis (see Section 2.1).

After preprocessing, artifact-free data were analyzed with Dynamic Causal Modelling (DCM), as this approach is highly indicated when CS presentation is longer than 0.75 seconds (Bach et al., 2010). To ensure accurate estimation of conditioned and unconditioned fear responses, only trials with less than a total of 6 seconds of missing epochs (after data cleaning, see above) were included in the analysis. For this reason, some trials could not be estimated due to the presence of an excessive number of missing values. For both fear acquisition and extinction, we modelled the conditions “Context” (fixed event), “CS onset” (fixed event), “CS interval” (flexible event), and “US” (fixed event) (Bach et al., 2010). It is important to note that PsPM assumes the event sequence to be the same for all trials. Hence, in the case of non-reinforced trials (no US), we modelled US omissions by adding an event of the same duration as the actual US in the place where the shock would have occurred during reinforced trials. As it is possible that the response to the CS+ is affected by the response to the US (due to overlap in their timings at the end), we applied two additional steps. First, we gradually decreased the time interval between CS+ onset and US onset to the point that there would be no differences between reinforced and non-reinforced CS+ trials. Second, for each trial, we ensured that the onset of the Gaussian bump plus at least two standard deviations did not overlap with the onset of the US. Both procedures ensured that the response to the CS+ during reinforced trials was not contaminated by the response to the US. Indeed, we found the difference between reinforced and non-reinforced CS+ trials not to be significant (*p* > .05), suggesting that our method appropriately disambiguated both types of response. SCR data obtained during fear acquisition and extinction training were normalized in the same way and analyzed via the same DCM approach to achieve a high degree of comparability. Once modelling was completed, the results of each trial were manually inspected to ensure a proper model fit. Finally, for each participant, we extracted the amplitude for each trial and for each phase of the experiment. These data, referred to as SCR values in the following, were subjected to statistical analysis.

### 2.4 Acquisition of Imaging Data

All imaging data were acquired at Bergmannsheil hospital in Bochum (Germany) using a 3 Tesla Philips Achieva scanner with a 32-channel head coil. Scanning included T1-weighted imaging, diffusion-weighted imaging, and resting-state imaging. The T1-weighted and diffusion-weighted images were obtained on day 1, whereas the resting-state images were obtained directly before the fear acquisition and fear extinction paradigm on day 2 (Figure 1).

### 2.5 T1-weighted Imaging

For the purpose of brain image parcellation, and in order to quantify macrostructural parameters such as regional brain volume (VOL), cortical thickness (THIC), and cortical surface area (SURF), T1-weighted high-resolution anatomical images were acquired (MP-RAGE, TR = 8.2 ms, TE = 3.7 ms, flip angle = 8°, 220 slices, matrix size = 240 mm x 240 mm, resolution = 1 mm x 1 mm x 1 mm). Scanning time was around 6 minutes.

#### 2.5.1 Preprocessing of T1-weighted Data

For the purpose of reconstructing the cortical surfaces of T1-weighted images, we used published surface-based methods in FreeSurfer (http://surfer.nmr.mgh.harvard.edu, version 6.0.0) along with the CBRAIN platform (Sherif et al., 2014). The details of this procedure have been described elsewhere (Dale et al., 1999; Fischl et al., 1999). The automatic reconstruction steps included skull stripping, gray and white matter segmentation, as well as reconstruction and inflation of the cortical surface. These preprocessing steps were performed for each participant individually. Subsequently, each individual segmentation was quality-controlled slice by slice, and inaccuracies of the automatic steps were corrected by manual editing if necessary.

The analyses of structural and functional brain connectivity, macrostructural brain properties, and NODDI coefficients followed a region-based approach, for which we considered brain regions as defined by the Human Connectome Project’s multi-modal parcellation (HCPMMP; the complete list of brain regions adapted from Glasser et al., 2016 is presented in Supplementary Table S7) and FreeSurfer’s automatic subcortical segmentation (Figure 1). The HCPMMP delineates 180 cortical brain regions per hemisphere and is based on the cortical architecture, function, connectivity, and topography of 210 healthy individuals. From FreeSurfer’s automatic subcortical segmentation, we included a set of eight subcortical regions per hemisphere (thalamus, caudate nucleus, putamen, pallidum, hippocampus, amygdala, insula, and accumbens area), various ventricle masks (lateral ventricle, inferior lateral ventricle, third ventricle, fourth ventricle), and a mask covering the whole cerebral white matter compartment (Fischl et al., 2002). In total, 360 cortical masks, 16 subcortical masks, 8 ventricle masks, and one white matter mask were selected and linearly transformed into the native spaces of the diffusion-weighted and resting-state images to guide further analyses. Ventricle and white matter masks were used exclusively for nuisance signal extraction and were not included as predictors in the regression models.

### 2.6 Diffusion-weighted Imaging

For fiber tracking and the analysis of NODDI coefficients, diffusion-weighted images were acquired using echo planar imaging (TR = 7652 ms, TE = 87 ms, flip angle = 90°, 60 slices, matrix size = 112 x 112, voxel size = 2 mm x 2 mm x 2 mm). Diffusion weighting was based on a multi-shell, high angular resolution scheme consisting of diffusion-weighted images for b-values of 1000, 1800, and 2500 s/mm^2^, respectively, applied along 20, 40, and 60 uniformly distributed directions. All diffusion directions within and between shells were generated orthogonally to each other using the MASSIVE toolbox (Froeling et al., 2016). Additionally, eight volumes with no diffusion weighting (b = 0 s/mm²) were acquired as an anatomical reference for motion correction and computation of NODDI coefficients. Diffusion-weighted data were collected with reversed phase-encode directions, resulting in pairs of images with distortions going in opposite directions. Scanning time was around 18 minutes.

#### 2.6.1 Preprocessing of Diffusion-weighted Data

Diffusion-weighted data were prepared for fiber tracking and the computation of NODDI coefficients via a preprocessing pipeline comprising the following steps. First, images were corrected for signal drift (Vos et al., 2017) using ExploreDTI (Leemans et al., 2009). Second, we utilized the topup command from the FSL toolbox (Smith et al., 2004) to estimate the susceptibility-induced off-resonance field based on pairs of images with opposite phase-encode directions. Third, the topup output was used in combination with the eddy command (Andersson & Sotiropoulos, 2016), which is also part of the FSL toolbox, to correct for susceptibility, eddy currents, and head movement. Importantly, we also performed outlier detection during this step to identify slices where signal had been lost due to head movement during the diffusion encoding (Andersson et al., 2016).

Following preprocessing, the 376 cortical and subcortical regions (see Section 2.5.1) were transformed into native diffusion space and used as seed and target masks for probabilistic fiber tractography. We employed a dual-fiber model as implemented in the latest version of BEDPOSTX, which allows for the representation of two fiber orientations per voxel when more than one orientation is supported by the data. This enables modelling of crossing fibers and produces more reliable results compared with single-fiber models (Behrens et al., 2007). Probabilistic fiber tractography was carried out using the classification targets approach implemented in FMRIB’s Diffusion Toolbox (Behrens et al., 2003; Cohen et al., 2009). At each voxel, 5,000 tract-following samples were generated with a step length of 0.5 mm and a curvature threshold of 0.2 (only allowing for angles larger than 80°). The connectivity between a seed voxel and a specific target region was quantified as the number of streamlines originating in the seed voxel and reaching the respective target region, and the connectivity between two brain regions was computed as the sum of streamlines proceeding in both directions. By doing so, we obtained a structural connectome for each participant, comprising 376 network nodes connected via 70,500 unique edges, represented as a 376 × 376 connectivity matrix. From this matrix, we computed nodal efficiency scores (Wang et al., 2017) as implemented in the Brain Connectivity Toolbox (BCT; Rubinov & Sporns, 2010). This metric quantifies a region’s capacity for information transfer as the average inverse shortest path length between a node and all others in the network, with shortest paths identified via Dijkstra’s algorithm. This procedure yielded 376 nodal efficiency scores for diffusion-weighted imaging connectivity (DWINE) per participant, which were subsequently used as input features for ElasticNet fitting.

In addition to structural connectivity, we also extracted microstructural tissue properties from the same preprocessed diffusion-weighted data via neurite orientation dispersion and density imaging (NODDI; Zhang et al., 2012). NODDI coefficients were computed using the AMICO toolbox (Daducci, Canales-Rodriguez, et al., 2015), which reformulates the standard non-linear NODDI fitting as a convex optimization problem, dramatically reducing processing time (Daducci, Dal Palu, et al., 2015; Sepehrband et al., 2016). The NODDI technique features a three-compartment model distinguishing intra-neurite, extra-neurite, and CSF environments. For this study, we focused exclusively on the intra- and extra-neurite compartments. Intra-neurite environments are characterized by a stick-like, cylindrically symmetric diffusion signal arising from water molecules restricted by neurite membranes. In white matter, this reflects axonal geometry, while in gray matter, it serves as an indicator of the neuropil formed by dendrites and axons. For each voxel, NODDI estimates the intra-neurite volume fraction (INVF), i.e., the proportion of the diffusion signal attributable to intra-neurite environments (Jespersen et al., 2009, 2011; Zhang et al., 2012), which serves as a measure of neurite density. NODDI also provides the neurite orientation dispersion index (ODI), a tortuosity measure coupling the intra- and extra-neurite spaces. It reflects the alignment or angular dispersion of axons in white matter, or of axons and dendrites in gray matter (Billiet et al., 2015; Zhang et al., 2012). For each participant, mean values of INVF and ODI were computed across all 376 cortical and subcortical regions (see Section 2.5.1).

### 2.7 Resting-state Imaging

For the analysis of functional brain connectivity, fMRI resting-state images were acquired prior to fear acquisition training using echo planar imaging (TR = 2500 ms, TE = 30 ms, flip angle = 90°, 40 slices, matrix size = 112 mm x 112 mm, resolution = 2 mm x 2 mm x 3 mm). Participants were instructed to close their eyes, relax, and think of nothing in particular. Scanning time was around 8 minutes.

#### 2.7.1 Preprocessing of Resting-state Data

Resting-state data were preprocessed using MELODIC, which is part of the FSL toolbox. The preprocessing involved several steps: Discarding the first two volumes from each resting-state scan to allow for signal equilibration, motion and slice-timing correction, and high-pass temporal frequency filtering (0.005 Hz). Spatial smoothing was not applied in order to avoid the introduction of spurious correlations in neighboring voxels.

For each cortical and subcortical region (see Section 2.5.1 Preprocessing of T1-weighted Data), 376 in total, we calculated a mean resting-state time course by averaging the preprocessed time courses of corresponding voxels. Next, we computed partial correlations between the average time courses of all pairs of brain regions, while controlling for several nuisance variables. We regressed out the trajectories of 6 head motion parameters as well as the mean time courses extracted from the white matter and ventricle masks (Fraenz et al., 2021). The resulting partial correlation coefficients were subjected to a Fisher z-transformation (Fisher, 1921), which produced normally distributed data suitable for further testing. To reduce spurious connections, we pruned each matrix using the Orthogonal Minimum Spanning Trees (OMST) approach (Dimitriadis et al., 2017a, 2017b). This data-driven method avoids arbitrary thresholding by iteratively aggregating minimum spanning trees until an optimal balance between global efficiency and wiring cost is achieved, resulting in a sparse yet fully connected graph for every participant. Resting state nodal efficiency (RSNE) was then computed on these pruned graphs using the same BCT implementation described above, again yielding 376 efficiency scores per participant for ElasticNet fitting.

### 2.8 Statistical Analyses

All statistical analyses were conducted in either MATLAB (version R2022b) or Python. Regularized regression was performed using the scikit-learn package (Pedregosa et al., 2011), while ANOVA and t-tests of SCRs were carried out using the Pingouin (Vallat, 2018) and Scipy Python packages.

To quantify fear acquisition, we computed the difference between mean SCR amplitudes to CS+ and CS− trials. A larger difference reflects greater physiological responding to the CS+ compared to the CS−. Where possible, all 16 CS+ and 16 CS− trials from fear acquisition training were averaged to derive each participant’s mean SCR. In cases where individual trials were too noisy for inclusion, the remaining valid trials were used instead. The overall effect of fear acquisition was assessed using a one-way repeated-measures ANOVA with CS type (CS+ vs. CS−) as the within-subjects factor, examining whether mean SCR amplitudes differed significantly between CS+ and CS− across participants.

Fear extinction SCRs is typically characterized by elevated CS+ responses during early trials that diminish over time, while CS− responses remain relatively stable and slowly habituate. To capture this dynamic, we quantified extinction as a difference-of-differences score: we first computed the CS+ - CS− difference separately for the early and late halves of extinction (each based on four trials), then subtracted the late-half difference from the early-half difference. A strong extinction effect is reflected by a large early CS+ - CS− difference that substantially decreases, approaching zero by the late half. No trials required omission in this phase (see Section 2.2). This interaction between stimulus type (CS+ vs. CS−) and time (early vs. late extinction) was evaluated using a two-way repeated-measures ANOVA.

### 2.9 Bootstrapped and Cross-Validated ElasticNet Regression for Neural Predictors

To examine relationships between neural measures and individual differences in fear acquisition or extinction, we used ElasticNet regression via the ElasticNetCV function in scikit-learn. Unlike conventional regression, this regularized approach does not rely on p-values, thereby circumventing the need for multiple comparison corrections such as FDR or Bonferroni (Gongora et al., 2020). Instead, regularization applies a penalty that shrinks small regression weights toward zero, retaining only robust, non-negligible effects. The penalty magnitude is determined through k-fold cross-validation to guard against overfitting, where the data are partitioned into k subsets, each serving once as a held-out test set while the remaining subsets form the training set.

ElasticNet regression combines lasso and ridge penalties (Zou & Hastie, 2005). Lasso regression can shrink coefficients exactly to zero and is well-suited when many predictors have negligible effects, whereas ridge regression only asymptotically approaches zero and is preferable when most predictors contribute meaningfully. ElasticNet offers a principled compromise when prior expectations about predictor relevance are limited, and it additionally shrinks the correlated predictors as a group, rather than randomly shrinking correlated predictors, in the case of lasso regression (Zou & Hastie, 2005). This approach has been successfully applied in related neuroimaging contexts, including studies linking white-matter microstructure to intelligence (Gongora et al., 2020) and polygenic scores to brain connectivity (Engler et al., 2025; Genç et al., 2023) as well as microstructure (Stammen et al., 2025). For more methodological details, please see Zou and Hastie (2005) and Serang et al. (2017).

#### 2.9.1 First-Level Modality-Specific Analysis

To identify robust neural predictors within each imaging modality, we embedded the cross-validated ElasticNet algorithm within a bootstrap resampling procedure. For each modality, fear acquisition or extinction SCRs served as the dependent variable, with one of seven brain parameters (VOL = regional brain volume; SURF = cortical surface area; THIC = cortical thickness; INVF = intra-neurite volume fraction; ODI = orientation dispersion index; RSNE = resting-state connectivity nodal efficiency; DWINE = diffusion-weighted imaging connectivity nodal efficiency) constituting the independent variables. Across 1,000 bootstrap iterations, samples of the same size as the original dataset were drawn with replacement, with out-of-bag (OOB) samples reserved as held-out test sets to evaluate generalization performance.

To control for demographic confounds without introducing data leakage, age and sex were orthogonalized from all variables within each bootstrap iteration. Following the Frisch-Waugh-Lovell theorem (Frisch & Waugh, 1933; Lovell, 1963) and principles of debiased machine learning (Chernozhukov et al., 2018), separate linear regression models were fit on training data to predict both the target variable and each brain feature from age and sex. The resulting residuals were used for all subsequent modeling, with the same mapping functions applied to residualize the OOB test set. Residualized training data were then passed into an ElasticNet model with k=5 internal cross-validation folds. We recorded the selection frequency of each feature, defined as the proportion of iterations in which it received a non-zero coefficient, and retained only those brain regions selected in at least 50% of iterations, providing a stringent threshold to distinguish robust signals from sample-specific noise.

#### 2.9.2 The Second-Level Multimodal Stacked Analysis

To further refine the predictive model, we conducted a more conservative, exploratory second-level analysis in which the bootstrapped ElasticNet procedure from the first level served as an initial feature selection step (full results reported in Supplementary Materials). For each learning phase (fear acquisition and extinction training), stable brain features identified across all seven modality-specific analyses were concatenated into a single, comprehensive multimodal feature matrix. This combined matrix was then submitted to a second round of bootstrapped ElasticNet regression using 500 iterations, while preserving the identical methodological pipeline: fold-nested confound residualization via the Frisch-Waugh-Lovell approach, k=5 internal cross-validation folds, and L1 ratio grid search. Brain regions retaining a non-zero coefficient in at least 50% of second-level iterations are reported in the Supplementary Materials (Supplementary Figures S3-S7, and Supplementary Tables S3-S6). This stacked multimodal model represents the most conservative output of our analytical framework, isolating only those neural predictors that demonstrate unique, cross-modally robust explanatory power for individual differences in fear and extinction learning.

## 3. Results

### 3.1 Skin conductance responses

#### 3.1.1 Acquisition training

Mean SCR values from CS+ and CS− presentations during fear acquisition were compared using a one-way repeated-measures ANOVA examining the effect of CS type (CS+ vs. CS−) on average SCR across all trials (Figure 2, left). Results revealed a significant main effect of CS type, *F*(1, 100) = 10.0, *p* = .002, partial *η²* = 0.09, 90% CI = [0.02, 0.18], with SCRs significantly higher towards CS+ than CS−, indicating that fear acquisition was successful.

#### 3.1.2 Extinction training

The SCR values from fear extinction were also averaged across CS+ and CS− trials but separately for the early and late extinction halves (Figure 2, right). These mean values were subjected to a two-way repeated measures ANOVA. The two factors were “stimulus type” (CS+ vs. CS−) and “time” (early vs. late). We found a significant main effect of “stimulus type”, *F*(1, 87) = 4.20, *p* = .04, partial *η²* = 0.05, 90% CI = [0.00, 0.13], and a significant main effect of “time”, *F*(1, 87) = 8.42, *p* = .005, partial *η²* = 0.08, 90% CI = [0.02, 0.19]. Results also showed a significant interaction effect between “stimulus type” and “time” *F*(1, 87) = 5.45, *p* = .02, partial *η²* = 0.05, 90% CI = [0.01, 0.15], indicating that the difference between mean CS+ and CS− values had changed substantially from the early to the late half of fear extinction. The Bonferroni-corrected post hoc tests showed that SCRs were significantly higher for CS+ than CS– only in the early but not the late half: early CS+: *M* = 0.44, *SD* = 0.76; early CS−: *M* = 0.30, *SD* = 0.44, 95% CI [.04, .26]; *p* = .02, Cohen’s *d* = 0.23, late CS+: *M* = 0.23, *SD* = 0.29; late CS−: *M* = 0.23, *SD* = 0.38, 95% CI [-.08, .07]; *p* > 0.99, Cohen’s *d* < -0.01. To further examine the time course of extinction, SCRs were compared between the early and late halves of extinction separately for CS+ and CS−. CS+ responses decreased significantly from early (*M* = 0.44, *SD* = 0.76) to late extinction (*M* = 0.23, *SD* = 0.29; 95% CI = [0.06, 0.35], *p* = .009, Cohen’s *d* = 0.365), indicating successful within-session extinction of conditioned fear. No significant change was observed for CS− responses across halves (early: M = 0.30, SD = 0.44; late: *M* = 0.24, *SD* = 0.38, 95% CI = [-0.00, 0.13]; *p* = .167, Cohen’s *d* = 0.151).

**Figure 2.**
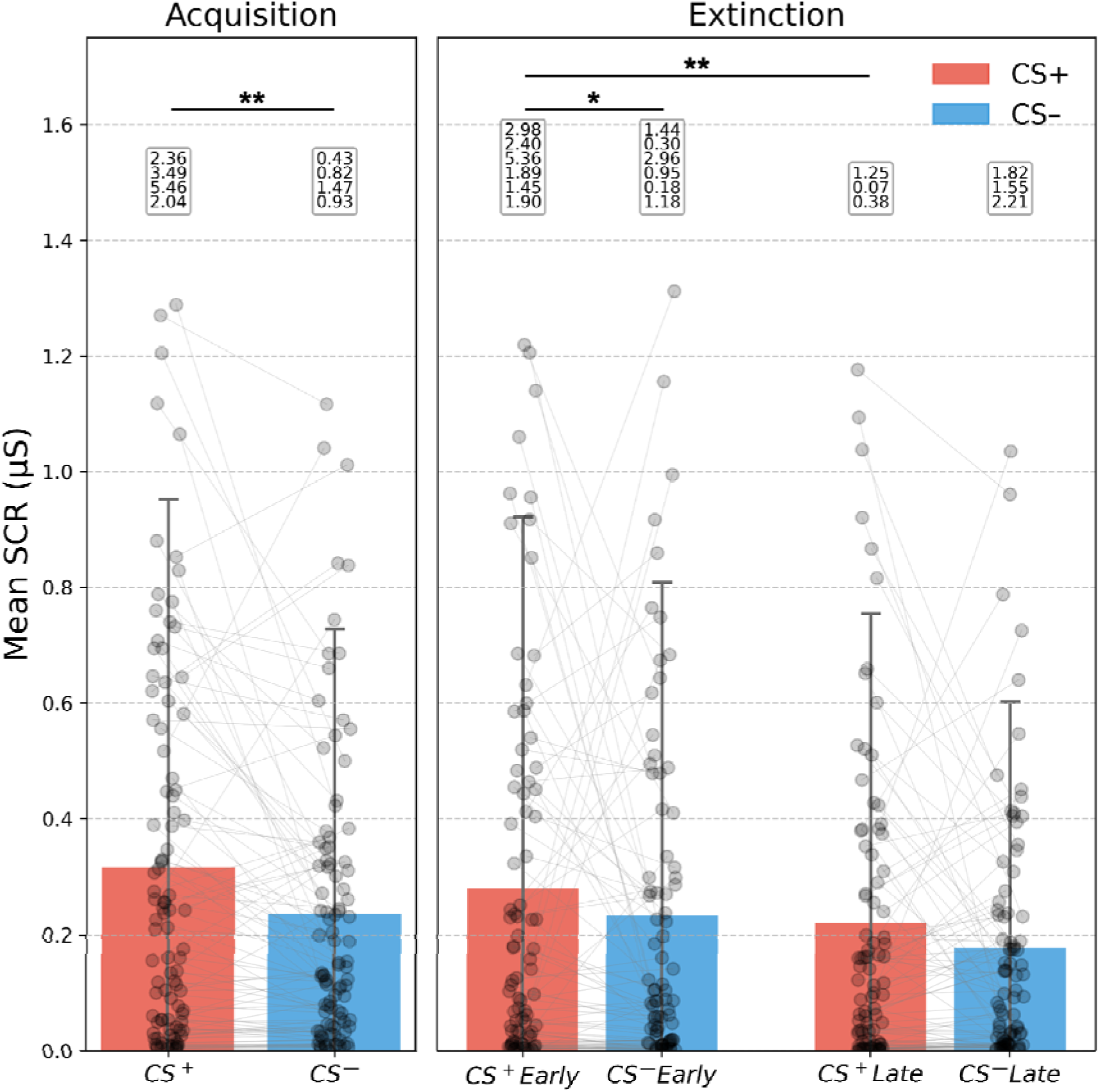
Mean skin conductance responses. Box plots showing the distribution of mean SCRs in reaction to CS+ (red boxes) and CS− (blue boxes) presentations during fear acquisition and extinction. In case of fear acquisition, the data were averaged across all 16 CS+ or 16 CS− presentations, respectively. In case of fear extinction, data were averaged across the first or last four CS+ or CS− presentations, respectively. For all plots, mean SCRs are given in microSiemens (µS). For each distribution, the whiskers represent 95% CI around the mean. Outliers are not shown visually but listed in white boxes for the sake of readability (** *p* < .01, * *p* < .05).

### 3.2 Modality-specific predictive markers

To identify the most reliable neural predictors of fear acquisition and extinction, we employed ElasticNet regression across seven distinct imaging modalities. This high-dimensional approach revealed a diverse set of regional parameters, spanning volume, surface area, cortical thickness, microstructure, and connectivity, that predicted differential SCRs in fear acquisition and a difference between early and later differential SCRs in extinction training, as well as the overlapping, dynamic brain areas that predicted both phases (see Figure 3 below; the complete list of per-modality brain areas can be found in Supplementary Figures S1 and S2 and Supplementary Tables S1 and S2). The OOB metrics of first-level and second-level models, as a result of bootstrapped, nested cross-validation, are described in Supplementary Tables S3 and S4.

**Figure 3.**
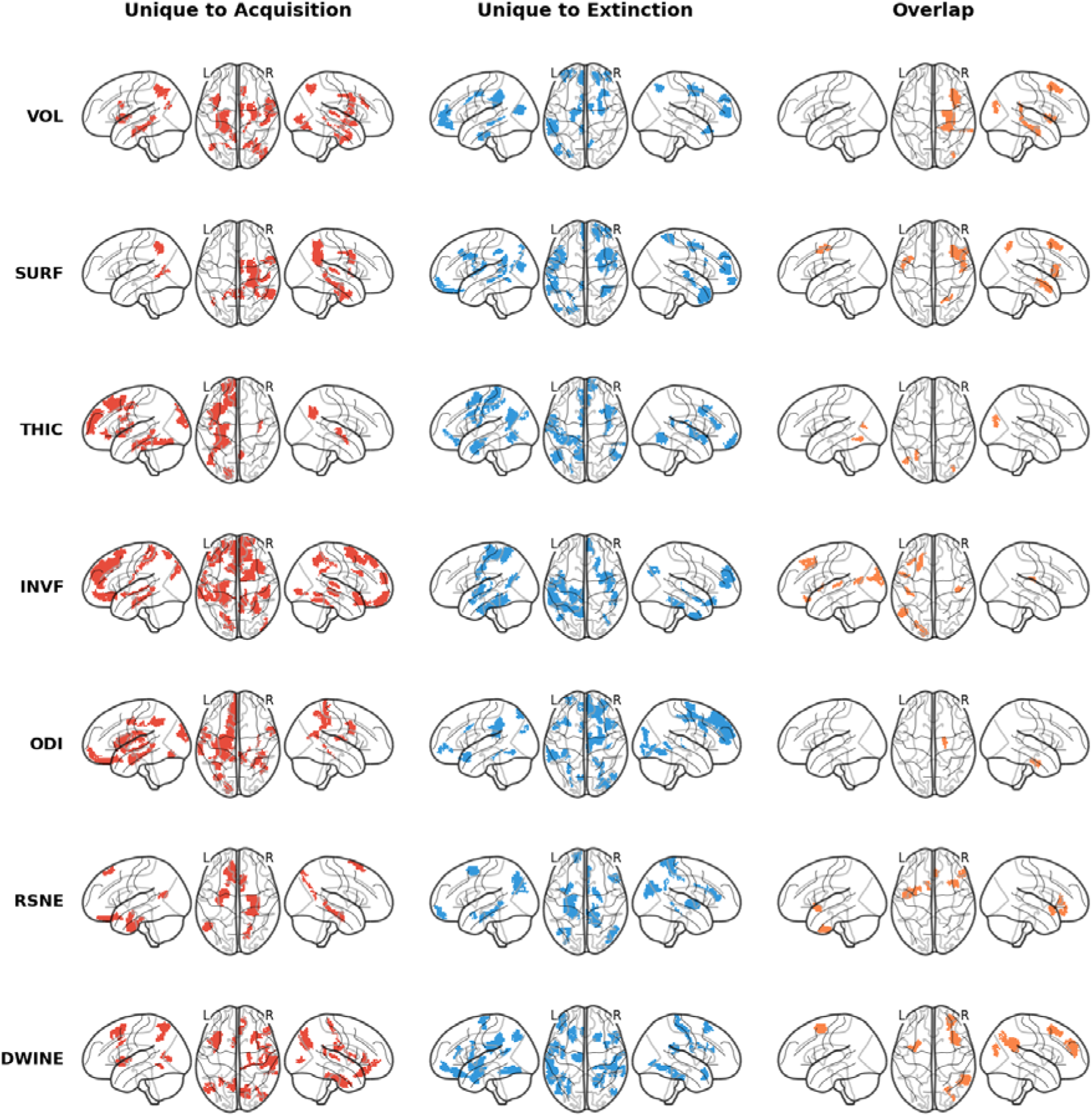
Modality-specific brain regions predictive of physiological fear responses across seven neuroimaging modalities. Results represent brain regions that were consistently assigned a non-zero weight in at least 50% of 1,000 bootstrap iterations of nested cross-validated ElasticNet regression at the first level (i.e., each modality fitted independently). Each row corresponds to one imaging modality (VOL = regional brain volume; SURF = cortical surface area; THIC = cortical thickness; INVF = intra-neurite volume fraction; ODI = orientation dispersion index; RSNE = resting-state connectivity nodal efficiency; DWINE = diffusion-weighted imaging connectivity nodal efficiency). Columns display regions consistently predictive of differential SCRs during fear acquisition (left, red), regions consistently predictive of the difference of the differential SCRs over the course of fear extinction (middle, blue), and the overlapping regions that were predictive during both phases (right, orange). Glass brain views show left lateral, dorsal, and right lateral perspectives for each condition. The complete list of brain areas and their bootstrap stability statistics is provided in Supplementary Tables S1 and S2, and visualized in Supplementary Figures S1 and S2.

#### 3.2.1 Fear acquisition training

A widespread predictive network characterized fear acquisition. We found 21 brain regions that consistently had non-zero weights across more than 50% of 1,000 bootstrap iterations for cortical volume of a region (VOL), which spanned the classical fear network regions of right anterior cingulate cortex (R_a32pr and R_p24pr), left insula and operculum (L_MI), pre-subiculum, right premotor (R_6v), bilateral visual areas (R_V8, L_PCV, R_LO2), and the left hippocampus.

Similarly, the cortical surface area (SURF) revealed 13 brain regions, including posterior cingulate and visual areas (R_PCV, L_PCV, L_ProS, L_MST), medial temporal structures (R_H, R_PeEc, R_PreS), cingulo-premotor regions (R_p24pr, R_6v), and additional parietal, auditory, opercular, and inferior parietal areas (R_LIPd, R_LBelt, R_FOP1, R_PGi).

Cortical thickness (THIC) identified 18 acquisition-related regions and showed a marked left-lateralized pattern encompassing fronto-insular and inferior frontal cortices (L_FOP2, L_MI, L_IFJa, L_IFJp), dorsolateral and orbital-prefrontal areas (L_8BL, L_9a, L_8Ad, L_8Av, L_10d), premotor and visual regions (L_FEF, L_V6A, L_V3A, L_FFC), medial temporal structures (L_H, L_PHA3), the anterior cingulate cortex (L_a24), and the right posterior cingulate region R_7m.

The INVF modality yielded the broadest acquisition profile, with 32 regions spanning auditory cortices (L_A5, R_A1, R_STSdp, R_LBelt), dorsal and ventral visual areas (L_V6, L_V6A, R_LO2, R_VVC, R_VMV2), superior parietal cortex (L_VIP), extensive dorsolateral, orbital, and medial prefrontal/cingulate regions (e.g., L_10r, L_8BM, L_46, R_8BL, R_a24pr, R_10v), somatosensory and midcingulate regions, the left hippocampus, and the right anterior agranular insula.

The ODI modality identified 23 acquisition-specific regions distributed across auditory-insular territories (L_52, L_A5, R_Ig, L_Ig, L_43), premotor and midcingulate areas (R_6v, L_23c, L_24dv, R_5m), visual-parietal regions (L_V8, L_V3A, R_AIP, L_IP2, R_LIPv, L_LIPd, R_TPOJ3), medial temporal structures (L_H, R_PreS), orbitofrontal and medial prefrontal cortex (L_OFC, L_10v), as well as subcortical contributions from the left caudate and amygdala.

RSNE showed a more circumscribed acquisition pattern, comprising 8 regions centered on mnemonic-limbic structures (R_H, L_PeEc, left amygdala), prefrontal areas (L_8BL, R_SFL, L_OFC), and posterior association regions (L_TPOJ2, R_DVT).

Finally, DWINE revealed 24 acquisition-specific regions, including occipito-parietal and MT+ areas (R_LO1, L_VMV2, L_FST, L_MST, R_V6, R_POS2), fronto-insular and opercular regions (R_FOP3, L_MI, R_FOP1), inferior parietal cortex (R_IP0, R_PFop, L_LIPd), medial frontal areas (R_p32, R_s32, L_a24pr, L_8Av), auditory and inferior frontal regions (R_LBelt, R_45, R_47s), and medial/lateral temporal structures (R_EC, R_TE2a).

The complete list of per-modality brain areas can be found in Supplementary Figure S1 and Supplementary Table S1, and is visualized in Figure 3 below (left/red).

#### 3.2.2 Extinction training

Predictors of extinction were similarly distributed across modalities, although the overall pattern shifted toward frontoparietal, cingulo-opercular, and higher-order sensory association regions.

For SURF, 23 regions were identified, with prominent superior parietal and posterior visual areas (R_7PC, L_V3B, L_DVT, L_VMV2, L_LO3), dorsolateral and orbital-prefrontal cortex (R_a9-46v, L_8C, R_a10p, R_9p), inferior frontal and opercular cortices (R_IFJa, L_IFJp, L_43, L_OP4, L_FOP1), premotor regions (R_FEF, R_6a), and temporo-occipital and auditory association areas (L_PSL, L_TPOJ1, L_STSvp, R_TGd).

THIC identified 21 extinction-specific regions, emphasizing midcingulate and paracentral territories (L_SCEF, L_5mv, R_p24, L_p32), somatosensory and premotor cortex (L_1, L_3a, L_6v), insular and opercular regions (R_PoI2, R_OP2-3, R_AAIC), dorsal visual and MT+ areas (L_V7, L_PH, R_PH), lateral temporal cortex (L_TE1a), and anterior cingulate/orbitofrontal areas (L_s32, R_10pp, L_a24pr).

The INVF modality revealed 20 extinction-related regions spanning bilateral early auditory cortex (L_52, R_52), somatomotor and parietal cortex (L_4, L_7PC, L_7AL, R_IP0), posterior and para-insular regions (L_PoI2, L_PI, R_PI), lateral temporal cortex (L_TE2a, L_TE2p, R_TE2p, R_TGv), medial temporal cortex (L_PHA1), temporo-parieto-occipital association cortex (L_TPOJ3), and frontal-orbitomedial areas (R_9m, R_13l).

ODI produced the largest extinction microstructural pattern, with 25 regions covering early and ventral visual areas (R_V4, L_VMV3, R_VMV3, R_V4t, L_V7), fronto-cingulate and dorsolateral prefrontal cortex (R_8BL, R_8C, R_9-46d, R_8Ad, R_9m, L_p32, L_a24pr), inferior frontal and premotor cortices (R_IFJa, R_6a), parietal regions (L_PF, L_7PL), medial temporal cortex (L_PHA2), orbitofrontal cortex (L_a10p, L_47s), and auditory association areas (R_52, L_RI, L_STGa).

RSNE yielded 19 extinction-specific regions centered on medial temporal structures, bilateral pre-subiculum, left hippocampus (L_PreS, L_H, R_PreS), inferior frontal cortex (R_IFSp), superior and inferior parietal regions (R_7AL, L_IP1, L_IP0, R_PGp), midcingulate and posterior cingulate areas (R_5L, R_5m, R_5mv, R_p24, R_33pr, R_31pd), orbitofrontal cortex (L_10pp), piriform cortex, and the temporo-parieto-occipital junction (L_TPOJ3).

DWINE identified 25 extinction-specific regions, highlighting inferior frontal regions (R_IFSp, R_IFJp, L_a47r), inferior parietal cortex (L_PF, L_IP1, L_PFop), ventral visual and fusiform areas (L_FFC, L_VMV3, R_VMV1, R_VMV3, R_V4t), medial and lateral temporal cortex (R_PHA3, R_TE2p, L_TE1a), cingulate and premotor regions (L_p24, L_6r, L_6mp), auditory cortex (R_STGa, L_TA2), and higher-order frontal areas (R_s6-8, L_13l).

The complete list of per-modality brain areas can be found in Supplementary Figure S2 and Supplementary Table S2 and is visualized in Figure 3 below (middle/blue).

#### 3.2.3 Overlapping predictors between acquisition and extinction

A subset of regions demonstrated dynamic predictive power, predicting differential SCRs during fear acquisition training as well as the difference in differential SCRs over the course of extinction training.

In VOL, this overlap was restricted to five regions spanning the right hippocampus, right middle insula, dorsolateral prefrontal cortex (R_8Av), a visual association area (R_V3CD), and superior temporal visual cortex (R_STV).

SURF also showed five overlapping regions, linking medial intraparietal and premotor cortex (R_MIP, L_55b) with auditory and inferior frontal areas (R_STSda, R_44), and again the right dorsolateral prefrontal cortex (R_8Av).

For THIC, dynamic effects were concentrated in three visual regions (L_VMV3, L_MT, R_V3CD), whereas INVF showed a broader overlap of eight areas encompassing orbitofrontal and dorsolateral prefrontal cortex (L_47m, L_8Ad), temporo-parieto-occipital association cortex (L_TPOJ2), dorsal visual cortex (L_V3B, L_V3A), primary auditory cortex (L_A1), and posterior opercular regions (L_FOP1, R_OP1).

Dynamic overlap was comparatively sparse for ODI, which retained only the right entorhinal cortex, while RSNE showed five overlapping regions, including the accumbens, right s32, right anterior agranular insula, right area 45, and left TGv.

In DWINE, six regions remained predictive across both phases, spanning visual and parietal cortex (R_V3CD, R_IPS1, R_PGi) and dorsolateral prefrontal areas (R_9-46d, R_8Av, L_i6-8).

### 3.3 Consensus of regions converging across modalities

To distill the most robust neural markers, we next identified regions that recurred across multiple imaging modalities, thereby providing a consensus view of the neural architecture supporting fear acquisition and extinction (Figure 4 left/red and middle/blue). We further examined regions that showed cross-phase overlap, that is, areas whose predictive value generalized across both acquisition and extinction despite differences in modality (Figure 4, right/orange).

**Figure 4.**
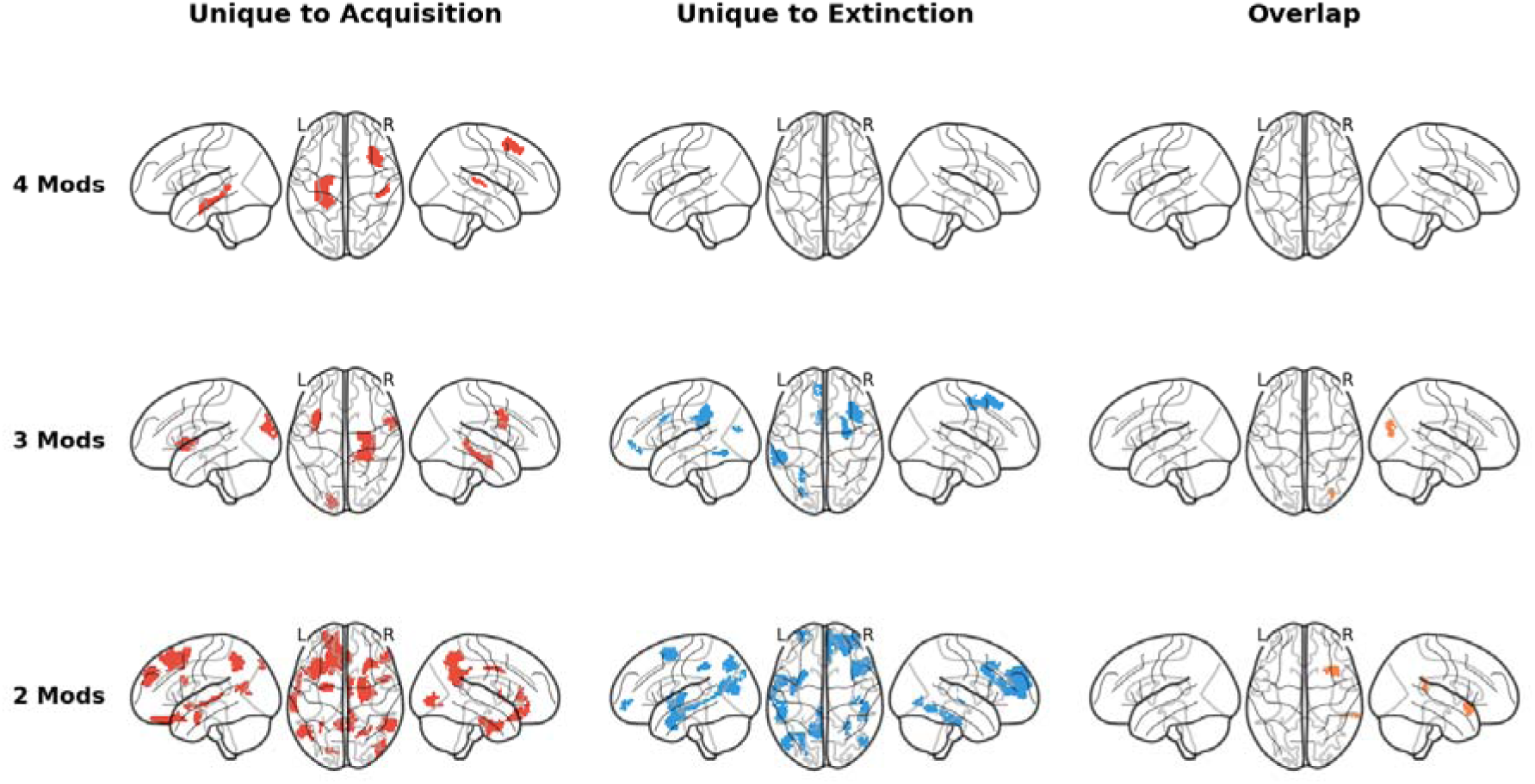
Brain regions whose predictive relationship with fear acquisition and extinction replicates across multiple neuroimaging modalities. To identify regions whose association with conditioned fear responses is not modality-specific, we examined which areas were selected by ElasticNet regression across two or more of the seven imaging modalities (first-level analysis; ≥50% bootstrap stability per modality). Rows indicate the level of cross-modal consensus, showing regions identified in at least four (top), three (middle), or two (bottom) distinct modalities. The color convention follows that of Figure 3: red = fear acquisition predictors; blue = fear extinction predictors; orange = dynamic predictors (predictive in both phases). Columns display regions consistently predictive of differential SCRs during fear acquisition (left, red), regions consistently predictive of the difference of the differential SCRs over the course of fear extinction (middle, blue), and the overlapping regions that were predictive during both phases (right, orange). No region reached consensus across all seven modalities. The complete list of brain areas and their bootstrap stability statistics is provided in Supplementary Tables S1 and S2, and visualized in Supplementary Figures S1 and S2.

#### 3.3.1 Fear acquisition training

At the highest overlap across modalities, three acquisition-specific regions were identified across four modalities: the left hippocampus (L_H), the right lateral belt complex (R_LBelt), and the right dorsolateral prefrontal cortex (R_8Av). When the threshold was lowered to three modalities, five additional acquisition-related regions emerged, including the right hippocampus (R_H), ventral premotor cortex (R_6v), the left middle insular area (L_MI), the right presubiculum (R_PreS), and the left dorsal visual area V3A (L_V3A). At a more liberal two-modality threshold, the acquisition consensus profile expanded further to include limbic and medial temporal regions such as the amygdala, perirhinal/ectorhinal cortex, and entorhinal cortex, together with orbitofrontal, cingulate, dorsolateral prefrontal, parietal, and occipito-temporal visual association areas.

#### 3.3.2 Extinction training

No extinction-specific region showed overlap across four modalities. However, seven regions overlapped across three modalities, comprising anterior cingulate and medial prefrontal areas (L_p32, L_a24pr), the right dorsolateral prefrontal cortex (R_8Av), right premotor cortex (R_6a), left inferior parietal cortex (L_PF), and visual association regions in both dorsal and ventral streams (L_V3B, L_VMV3). At the two-modality threshold, the extinction consensus map broadened to include additional inferior frontal, insular, lateral temporal, posterior cingulate, dorsolateral prefrontal, auditory, and visual regions, indicating a distributed but less strongly convergent cross-modality pattern than that observed for fear acquisition training.

#### 3.3.3 Overlapping predictors between fear acquisition and extinction

For regions shared across fear acquisition and extinction, no area showed an overlap across four modalities, and only one region survived overlap across three modalities: the right V3CD visual association area. At the two-modality threshold, two further dynamic regions were identified, namely the right anterior agranular insula complex (AAIC) and the right superior temporal visual area (STV).

### 3.4 Consensus across macrostructure, microstructure, and connectivity imaging modalities

To further simplify the multimodal findings, we grouped the imaging modalities into three broader domains: macrostructure (VOL, THIC, SURF), microstructure (INVF, ODI), and connectivity (RSNE, DWINE), and then identified brain regions that overlapped within each domain (see Figure 5, below). This approach allowed us to isolate convergent regional predictors within the structural scale, rather than across individual modalities alone.

**Figure 5.**
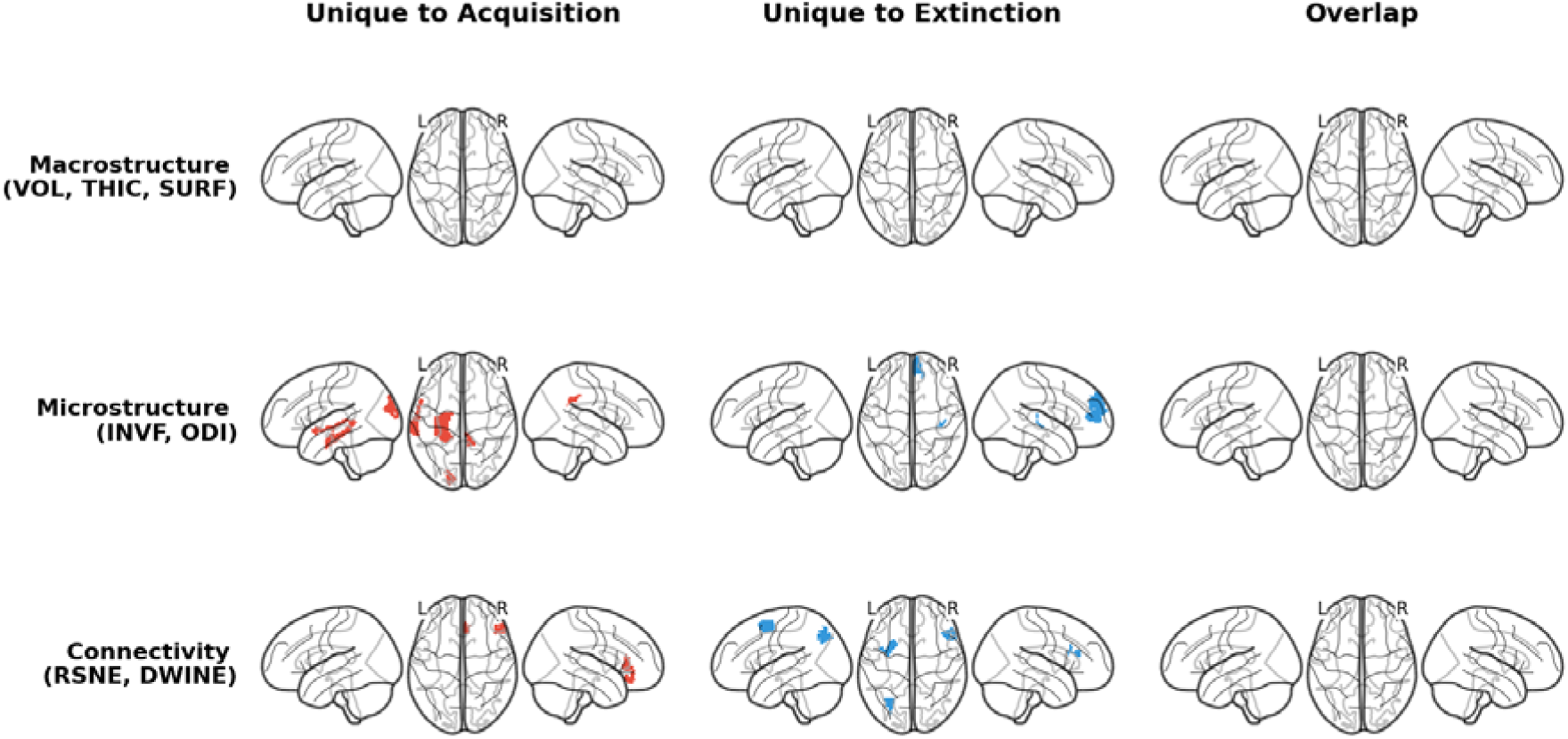
Convergence of fear learning predictors grouped across three neuroimaging domains: macrostructure, microstructure, and connectivity. To examine whether predictive brain regions generalize across methodologically distinct classes of imaging measure, modalities were grouped into three domains: macrostructure (VOL, SURF, THIC), microstructure (INVF, ODI), and connectivity (RSNE, DWINE). A region is shown if it was identified as a stable predictor (≥50% bootstrap stability) by all modalities, indicating convergent evidence that is unlikely to reflect modality-specific measurement variance. Rows represent the specific domain combinations showing overlap. Columns and color coding follow the convention of Figures 3 and 4 (red = fear acquisition; blue = extinction; orange = dynamic/overlap). Glass brain views show left lateral, dorsal, and right lateral perspectives. The full list of brain areas within each domain grouping is provided in Supplementary Figures S1 and S2.

#### 3.4.1 Fear acquisition training

For fear acquisition training, the macrostructural group did not yield any overlapping regions across VOL, THIC, and SURF. In contrast, the microstructural group showed a consensus of four regions, comprising the right posterior cingulate area 31pv (R_31pv), the left dorsal visual area V3A (L_V3A), the left auditory association area A5 (L_A5), and the left hippocampus (L_H). The connectivity group yielded two convergent acquisition-related regions, namely the right inferior frontal area 45 (R_45) and the right s32 region within anterior cingulate/medial prefrontal cortex (R_s32).

#### 3.4.2 Extinction training

A similar pattern was observed during extinction, where the macrostructural domain again showed no overlapping regions. The microstructural group identified two extinction-related consensus regions: the right early auditory area 52 (R_52) and the right area 9 middle (R_9m) within the anterior cingulate/medial prefrontal cortex. The connectivity domain showed a somewhat broader extinction consensus, including the right inferior frontal area IFSp (R_IFSp), the left inferior 6-8 transitional area (L_i6-8) within dorsolateral prefrontal cortex, and the left inferior parietal area IP1 (L_IP1).

#### 3.4.3 Overlapping predictors between fear acquisition and extinction

No regions showed overlap across both fear acquisition and extinction within any of the three broad modality domains. Thus, when modalities were collapsed into macrostructural, microstructural, and connectivity classes, cross-phase dynamic consensus was absent.

## 4. Discussion

In the present study, we examined how multimodal brain properties predict individual differences in fear and extinction learning as indexed by differential SCRs. Using bootstrapped, cross-validated ElasticNet models across seven imaging modalities, we identified modality-specific predictors of fear acquisition and extinction, regions showing convergence across modalities, and a set of areas with dynamic predictive value across both learning phases. Taken together, the findings indicate that fear conditioning overall is best understood as a distributed multimodal phenomenon, with fear acquisition showing broader and more robust convergence than extinction.

### Behavioral validation of fear acquisition and extinction

Before discussing the neuroimaging findings, it is important to note that fear acquisition and extinction training were successful on the behavioral level. During fear acquisition training, CS+ trials elicited significantly higher SCRs than CS− trials, confirming that participants successfully acquired conditioned fear responses, consistent with well-established differential conditioning outcomes (Lonsdorf et al., 2017; Milad, Wright, et al., 2007). During extinction training, CS+ responses were significantly elevated relative to CS− only in the early half of extinction but not in the late half, and CS+ responses decreased significantly from early to late trials, reflecting successful within-session extinction of conditioned responding (Lonsdorf & Merz, 2017). This interaction between stimulus type and time is the canonical hallmark of extinction learning (Bouton, 2004) and validates the SCR-based individual difference measures subsequently used as dependent variables in the neuroimaging analyses.

### Modality-specific predictors of fear acquisition

One of the key findings was that fear acquisition was predicted by a widespread set of cortical and subcortical markers across virtually all imaging modalities, with brain areas previously known to constitute the fear network. Across modalities, the acquisition pattern repeatedly involved hippocampal and parahippocampal regions, insular and cingulate cortex, dorsolateral and premotor frontal areas, as well as auditory and visual association cortices. This broad profile fits well with the idea that fear acquisition depends not on a single localized substrate, but on coordinated contributions from systems involved in salience detection, autonomic responding, contingency learning, memory formation, and sensory representation of threat-predictive cues (Milad & Quirk, 2012; Fullana et al., 2016). The VOL modality yielded 21 regions, including right anterior cingulate cortex, left insula and operculum, and left hippocampus, structures long implicated in fear and interoceptive processing (Buchel et al., 1999; Cauda et al., 2011). The cortical thickness modality showed a marked left-lateralized pattern encompassing fronto-insular, inferior frontal, dorsolateral, and orbital-prefrontal areas, consistent with previous findings relating cortical thickness of insular and cingulate cortex to conditioned fear responses (Hartley et al., 2011; Milad, Quirk, et al., 2007). The INVF modality yielded the broadest acquisition profile, with 32 regions spanning auditory, visual, prefrontal, and cingulate areas, suggesting that neurite density captures biologically relevant variance in these systems over and above macrostructural indices (Genç et al., 2018).

The hippocampal findings are especially consistent with a role in encoding and stabilizing CS/US contingencies. In fear conditioning, the organism must form a representation linking the conditioned cue to the aversive outcome and maintain that representation across repeated trials; multimodal predictive contributions from hippocampal and adjacent medial temporal structures fit well with this broader mnemonic function (Cacciaglia et al., 2015; Pohlack et al., 2012). The recurrent involvement of insular and cingulate regions is compatible with their established roles in salience processing, interoceptive awareness, and autonomic regulation, all of which are central to differential SCR expression during fear acquisition (Milad & Quirk, 2012; Apkarian et al., 2005). The repeated contribution of dorsolateral prefrontal regions across modalities also deserves emphasis. Rather than suggesting that fear acquisition is purely “emotional” or “limbic,” the present findings support a model in which executive and attentional control systems are already engaged during initial learning (Abend et al., 2020). One plausible interpretation is that dorsolateral and premotor regions contribute to maintaining task contingencies, allocating attention to predictive cues, and coordinating anticipatory control processes that shape autonomic differentiation between CS+ and CS−.

Another striking feature of the acquisition results was the robust involvement of sensory and association cortex, including visual, auditory, parietal, and temporo-occipital regions. This is unlikely to be explained solely by low-level perceptual demands, because the task-relevant stimulus discrimination itself was simple. Instead, the findings are more consistent with the view that sensory cortices participate directly in representing threat-predictive information (Wen et al., 2024), with higher-order sensory areas contributing to the perceptual encoding and updating of learned cue significance, a pattern also noted in recent large-scale MVPA work showing numerically superior decoding of conditioned threat using whole-brain patterns beyond the canonical “threat circuit” (Wen et al., 2024).

### Modality-specific predictors of fear extinction

In contrast to fear acquisition, extinction was characterized by a distributed but less strongly convergent predictive architecture. Although several modalities identified extinction-related markers in frontoparietal, cingulo-opercular, lateral temporal, parietal, and visual association regions, no region survived overlap across four modalities. This suggests that extinction learning is not absent at the neural level, but rather more heterogeneous and potentially more dependent on the specific brain property being measured. Fear extinction is widely understood to involve predominantly the formation of new inhibitory memories that compete with, rather than fully erase, the original threat association (Bouton, 2004; Milad & Quirk, 2012), although extinction-related weakening or updating of the original association may also contribute. This process may require flexible coordination between executive-control and association systems rather than the strongly unified multimodal signature characteristic of acquisition.

The frontoparietal and cingulo-opercular pattern observed during extinction is consistent with prior work implicating these systems in attentional control, performance monitoring, and the updating of learned associative value (Milad, Wright, et al., 2007; Fullana et al., 2018). Notably, the cortical thickness modality identified extinction-specific regions in midcingulate, somatosensory, premotor, insular, and medial prefrontal cortex, which broadly aligns with meta-analytic evidence for extinction-related activations in cingulate and prefrontal territories (Fullana et al., 2018). The SURF and DWINE modalities prominently featured inferior frontal and parietal cortex alongside dorsolateral prefrontal areas, consistent with the engagement of executive and attentional networks during the reduction of conditioned responding. This interpretation is further supported by recent large-scale neuroimaging work showing that fear extinction learning modulates functional connectivity across broadly distributed networks (including frontoparietal, default mode, and ventral attention systems) well beyond the canonical “fear circuit” (Wen et al., 2021), and that increases in large-scale connectivity during late extinction predict the magnitude of extinction memory recall (Wen et al., 2021).

### Overlapping dynamic predictors across both phases

A further important result was the relative sparsity of dynamic predictors shared across fear acquisition and extinction, similar to MVPA findings of Wen et al. (2024), which they referred to as “flexible dynamic coders”. Cross-phase overlap was limited overall, with only the right V3CD visual association area surviving consensus across three modalities, and only the right anterior agranular insular complex (AAIC) and the right superior temporal visual area (STV) emerged at the two-modality threshold. Rather than implying that fear acquisition and extinction are unrelated, this finding suggests that the two phases may recruit partially overlapping but largely non-identical predictive substrates, with only a narrow set of higher-order visual and insular-temporo-occipital regions showing consistent relevance across both phases. Fear acquisition and extinction differ substantially in their computational and psychological demands: fear acquisition requires building a threat prediction, whereas extinction requires updating that prediction in the absence of reinforcement (Bouton, 2004; Milad & Quirk, 2012). The present results fit the view that these phases share some perceptual- associative infrastructure yet differ in the broader constellation of neural properties that best predict individual differences in associative learning. Relatedly, Wen et al. (2024) showed using multivariate pattern analysis that many brain regions dynamically shift their representations between threat and safety across fear acquisition and extinction training, further supporting the notion that distinct neural substrates mediate different stages of fear and extinction learning.

### Consensus of regions converging across modalities

When we next identified regions that recurred across multiple imaging modalities, a consensus view of the neural architecture supporting each phase emerged. For fear acquisition, three regions emerged at the highest consensus level (four modalities): the left hippocampus, the right lateral belt complex, and the right dorsolateral prefrontal cortex (R_8Av). The hippocampal convergence across modalities is broadly consistent with prior structural studies linking hippocampal volume to fear conditioning outcomes (Cacciaglia et al., 2015; Pohlack et al., 2012; Winkelmann et al., 2016). The convergence of dorsolateral prefrontal cortex across macrostructural, microstructural, and connectivity modalities is similarly aligned with its established role in top-down regulation of conditioned responding (Abend et al., 2020; Milad, Quirk, et al., 2007). At the three-modality threshold, additional overlap included the right hippocampus, ventral premotor cortex, left middle insula, right presubiculum, and left V3A, pointing to a broader medial temporal–insular–prefrontal network as the shared multimodal substrate of fear acquisition. This pattern suggests that successful fear acquisition is characterized by jointly informative variation across memory-related medial temporal structures, salience-interoceptive regions, executive-prefrontal territories, and sensory-associative cortices, rather than by any single imaging modality in isolation (Wen et al., 2024; Fullana et al., 2016).

For extinction, no region showed an overlap across four modalities. Seven regions converged across three modalities: anterior cingulate and medial prefrontal areas (L_p32, L_a24pr), the right dorsolateral prefrontal cortex, right premotor cortex, left inferior parietal cortex, and visual association regions in both dorsal and ventral streams (L_V3B, L_VMV3). The prominence of medial PFC and anterior cingulate at the three-modality threshold is consistent with their well-documented roles in extinction memory formation and expression (Milad, Wright, et al., 2007; Sotres-Bayon et al., 2006), though previous work has predominantly characterized these regions through task-based fMRI activation rather than the structural and connectivity indices employed here. At the more liberal two-modality threshold, the extinction consensus map broadened to include additional inferior frontal, insular, lateral temporal, posterior cingulate, and auditory regions, indicating a distributed but less strongly convergent cross-modality pattern than that observed for fear acquisition. For cross-phase dynamic consensus, only the right V3CD visual association area survived three-modality overlap. At the two-modality threshold, the right AAIC and right STV emerged as additional dynamic consensus regions. Together, these findings indicate that cross-phase consensus was comparatively sparse and was centered primarily on higher-order visual and insular-temporo-occipital association cortex.

### Consensus across macrostructure, microstructure, and connectivity

This interpretation is reinforced by the domain-level analysis. When modalities were grouped into macrostructure (VOL, THIC, SURF), microstructure (INVF, ODI), and connectivity (RSNE, DWINE) classes, no overlapping macrostructural predictors emerged for either fear acquisition or extinction, whereas microstructural and connectivity markers showed modest but interpretable consensus. For fear acquisition, microstructural convergence involved the hippocampus, auditory cortex, posterior cingulate, and dorsal visual cortex, while connectivity convergence highlighted inferior frontal and medial prefrontal/cingulate regions. For extinction, microstructural overlap centered on the auditory and medial prefrontal cortex, and connectivity overlap involved inferior frontal, dorsolateral prefrontal, and inferior parietal regions. This pattern suggests that variation in tissue microstructure and network organization (as captured by NODDI-derived neurite density and tractography-derived connectivity) may provide more coherent predictive information for fear and extinction learning than gross morphometric measures alone (Genç et al., 2018; Nees et al., 2019; Fani et al., 2015). The absence of macrostructural consensus likely reflects the relatively coarser sensitivity of volumetric and cortical thickness measures compared with diffusion-based microstructure and functional connectivity, the latter being more responsive to subtle inter-individual variation in white matter architecture and network topology (Nees et al., 2019; Hermann et al., 2017). No regions showed cross-phase dynamic overlap within any of the three broad modality domains, further reinforcing the view that fear acquisition and extinction draw on largely distinct predictive substrates even when modalities are aggregated.

### Limited role of the amygdala

The amygdala played only a limited role in the present predictive models. Although the left amygdala appeared in some acquisition-related results (ODI and RSNE modalities), it did not emerge as a central multimodal consensus marker. This is noteworthy given the amygdala’s well-documented role in fear acquisition and expression (Buchel et al., 1999; LaBar et al., 1998; Phelps, 2004; Cacciaglia et al., 2015), but it is consistent with a growing body of evidence challenging the singular importance of amygdala responses in human fear conditioning (Visser et al., 2021; Fullana et al., 2019; Wen et al., 2022). In a whole-brain predictive framework using structural, microstructural, and connectivity features, the absence of strong amygdala convergence does not imply irrelevance; rather, it suggests that individual differences in SCRs were better captured by distributed cortical–medial temporal systems than by amygdala properties alone. This interpretation aligns with recent MVPA work demonstrating that whole-brain activation patterns beyond the canonical “threat circuit” numerically outperform amygdala-centered models in classifying conditioned threat from safety cues (Wen et al., 2024), and with evidence that the specific contributions of amygdala subregions to threat and safety learning are temporally and anatomically specific (Wen et al., 2022).

### Broader implications for a multimodal, network-based account of fear learning

More broadly, our findings argue for a holistic multimodal and network-based account of fear and extinction learning, rather than the single-modality perspective that has dominated the field. The strongest and most reproducible predictive patterns were not confined to canonical “fear regions,” but extended across memory, salience, executive, and sensory-association systems. This broader architecture is consistent with recent proposals that fear conditioning engages multiple functionally distinct neural networks, including those involved in attention, working memory, and conscious awareness, well beyond the traditional “threat circuit” (Wen et al., 2021; Wen et al., 2024; Fullana et al., 2016, 2018). Such distributed engagement may help explain why prior work focused on single modalities or a narrow set of regions has often produced fragmented findings. Different imaging features may capture complementary aspects of the same learning phenotype and allow better accounting of the interindividual variability (Gomes et al., 2026; Hartley et al., 2011; Lonsdorf & Merz, 2017).

### Limitations

One of the limitations of this study is that, due to the models being regularized and selection-based, the identified regions should be interpreted as potentially predictive markers rather than isolated causal loci. However, due to a small sample size (N=101 and N=88) relative to a high feature space (376 cortical or 360 subcortical regions), and despite the bootstrapping approach, which improves robustness, the precise regional composition of the predictive sets still requires replication in considerably larger samples, especially given the considerable p>>n problem (Vabalas et al., 2019) and the negative OOB R² metrics for the first-level models (Supplementary Tables S3 and S4). It is important to note that OOB R² metrics for a second-level, truly multimodal ElasticNet regression, with a shortlisted set of brain areas pulled across modalities from a first-level 1,000 iteration bootstrapped, cross-validated ElasticNet regression, point at a reliable and considerably higher predictive power of these metrics, which might have been limited by a small sample size and large feature space. Lastly, the broader acquisition profile relative to extinction may partly reflect the fact that acquisition produced a more robust and stable individual-difference signal in SCRs, whereas extinction depends on more fragile updating processes and may therefore yield weaker cross-modality agreement. This effect may also be partially explained by the fact that we used different brain properties to predict a difference of a difference of SCRs (learning rate) for extinction, which might be a more complex metric than a simple differential SCR, used in acquisition.

## 5. Conclusion

In conclusion, this study provides a more integrated account of the neurobiological architecture of fear and extinction learning. Fear acquisition showed broad and comparatively strong multimodal convergence, especially across hippocampal, insular, cingulate, dorsolateral prefrontal, and sensory-association regions, whereas extinction was predicted by a more heterogeneous frontoparietal and cingulo-opercular profile with weaker consensus. Overall, the results support the idea that individual differences in conditioned autonomic responding arise from distributed properties of brain structure and connectivity, and that multimodal prediction offers a useful framework for identifying neural markers of fear learning beyond any single modality or canonical region.

## Funding and Acknowledgements

This work is part of the Collaborative Research Center SFB 1280 on extinction learning (projects A02, A03, A09, and F02). It was supported by the Deutsche Forschungsgemeinschaft (DFG; project number 31680338). The authors would like to thank PHILIPS Germany (Burkhard Ma dler) for scientific support with the MRI measurements, as well as Tobias Otto for technical support. We would like to thank Carlos Alexandre Gomes for assistance with the analysis of SCR data in PsPM. The authors would also like to express their gratitude to all research assistants for their support with data acquisition.

## Author Contributions

**Arslan Gabdulkhakov:** Data Curation, Software, Investigation, Methodology, Formal Analysis, Writing - Original Draft, Writing - Review & Editing **Christoph Fraenz:** Formal analysis, Investigation, Data Curation, Writing - Original Draft, Writing - Review & Editing, Visualization; **Dorothea Metzen:** Formal analysis, Investigation, Data Curation, Writing - Original Draft, Writing - Review & Editing; **Julian Packheiser:** Writing - Original Draft, Writing - Review & Editing; **Christian J. Merz:** Methodology, Writing - Review & Editing; **Nikolai Axmacher:** Conceptualization, Writing - Review & Editing, Supervision, Project administration; **Erhan Genç:** Conceptualization, Methodology, Writing - Review & Editing, Supervision, Project administration

## Data Availability

Data and code used in the analyses are available from the corresponding author upon request.

