## Supplementary Materials for "Distinct and overlapping correlates of fear acquisition and extinction across different neuroimaging modalities"

**for**

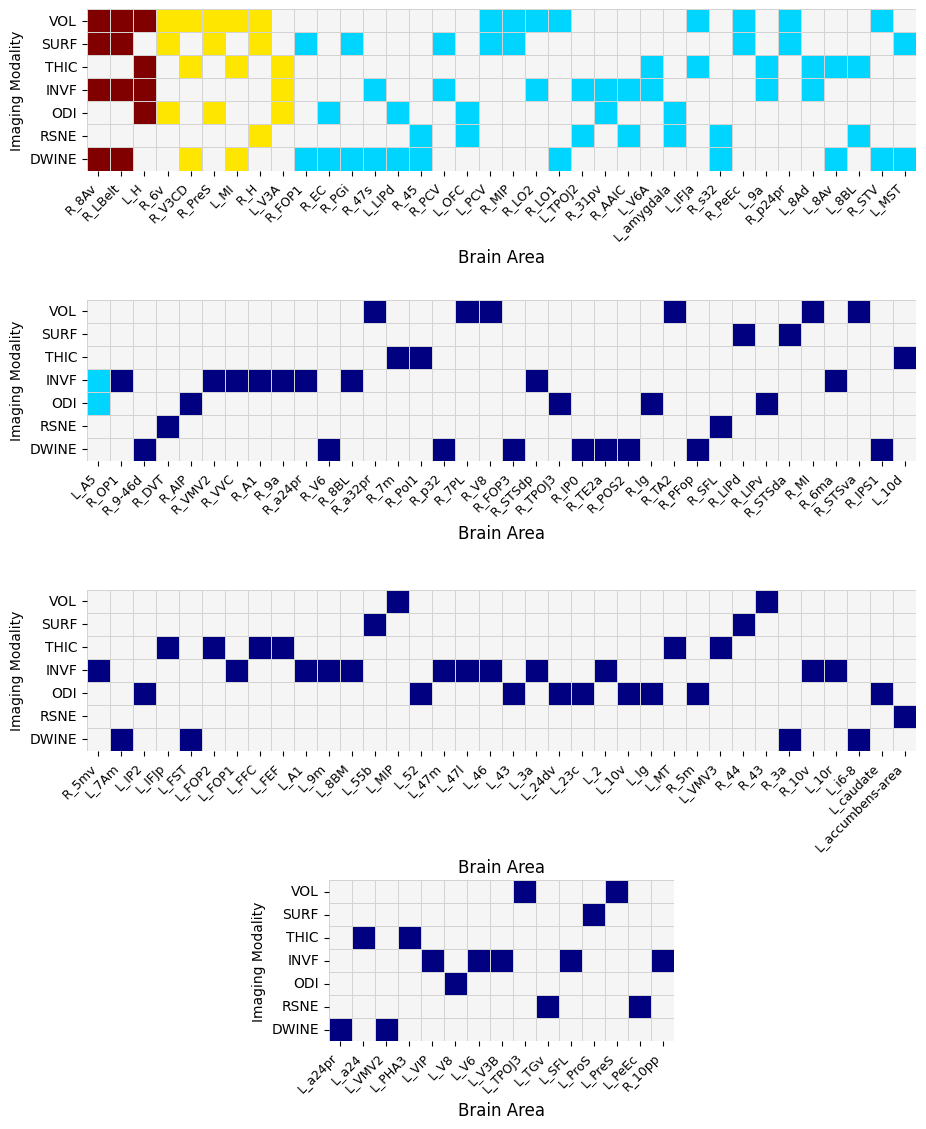

**Figure S1.** A schematic, visual depiction of the brain areas in each imaging modality that were consistently predictive of the SCR outcome of fear learning across at least 50% of 1000 bootstrap iterations of cross-validated ElasticNet. The **maroon**-filled cells represent the brain areas that were predictive of SCR-based learning outcome across 4 imaging modalities; **yellow**-filled cells represent the brain areas that were predictive of SCR-based learning outcome across 3 imaging modalities; **cyan**-filled cells represent the brain areas that were predictive of SCR-based learning outcome across 2 imaging modalities; **blue**-filled cells represent the brain areas that were predictive of SCR-based learning outcome in 1 imaging modality. The 'L_' and 'R_' prefixes to region codes correspond to the left hemisphere and the right hemisphere. The table with longer anatomical names of the corresponding coded areas is presented at the end of the supplementary materials, Table S7.

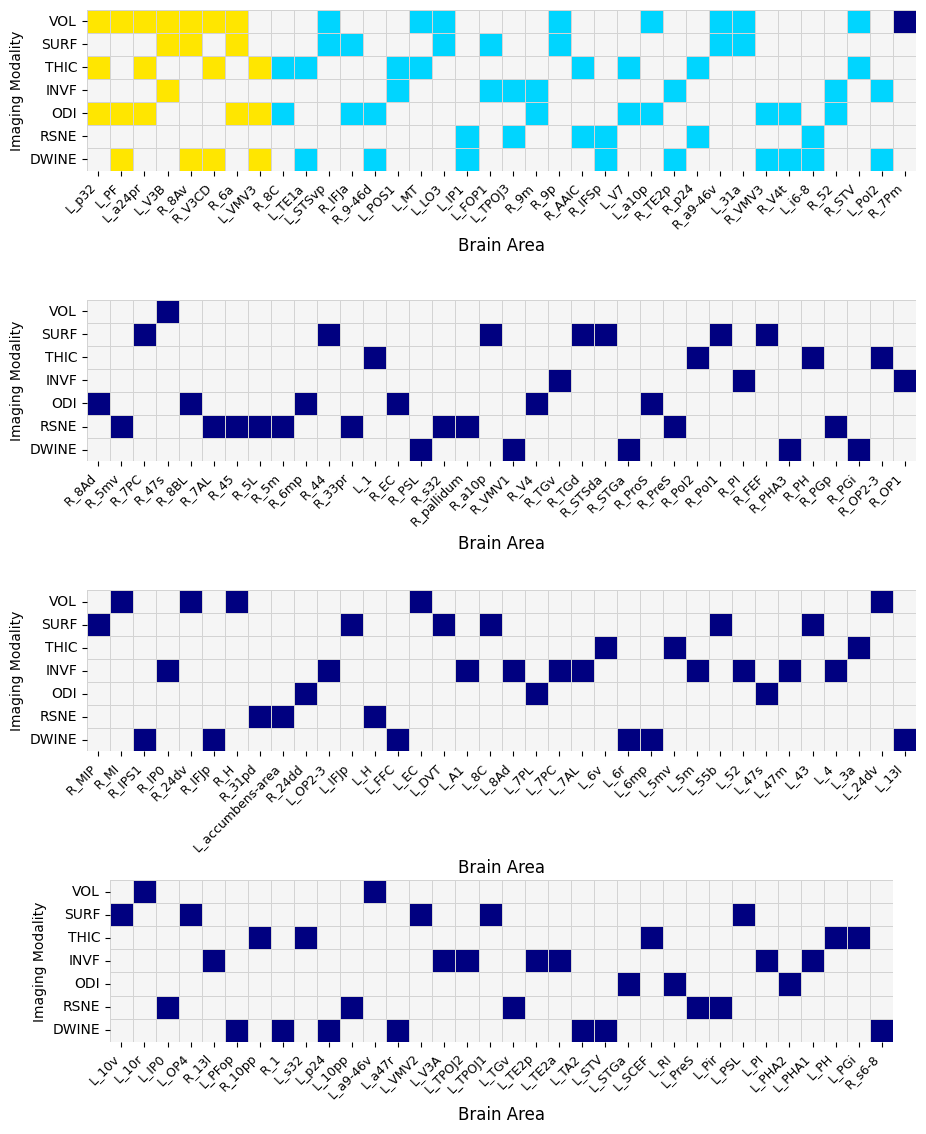

**Figure S2.** A schematic, visual depiction of the brain areas in each imaging modality that were consistently predictive of the difference of differential SCRs over the course of extinction learning across at least 50% of 1000 bootstrap iterations of cross-validated ElasticNet. The **maroon**-filled cells represent the brain areas that were predictive of SCR-based learning outcome across 4 imaging modalities; **yellow**-filled cells represent the brain areas that were predictive of SCR-based learning outcome across 3 imaging modalities; **cyan**-filled cells represent the brain areas that were predictive of SCR-based learning outcome across 2 imaging modalities; **blue**-filled cells represent the brain areas that were predictive of SCR-based learning outcome in 1 imaging modality. The 'L_' and 'R_' prefixes to region codes correspond to the left hemisphere and the right hemisphere. The table with longer anatomical names of the corresponding coded areas is presented at the end of the supplementary materials, Table S7.

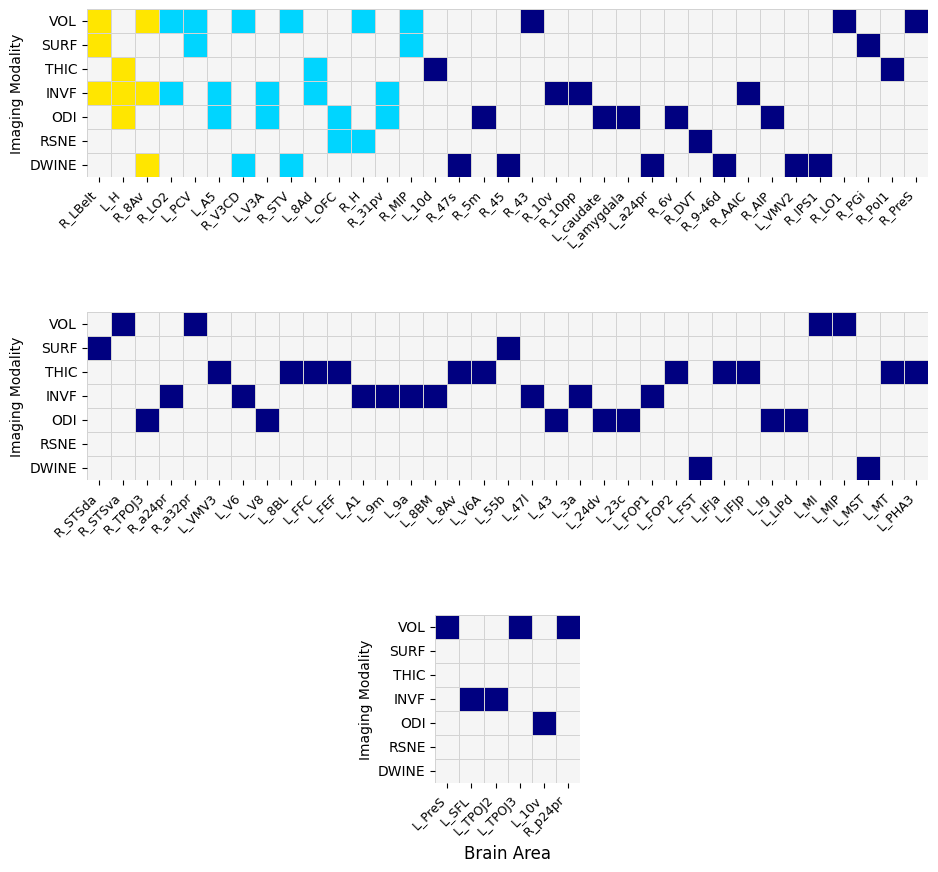

**Figure S3.** A schematic, visual depiction of the brain areas in each imaging modality that were consistently predictive of the SCR outcome of fear learning across at least 50% of 500 bootstrap iterations of cross-validated second-level multimodal ElasticNet, which received the shortlisted brain areas from all imaging modalities of an earlier 1000-iteration bootstrapped cross-validation in Figure S1. The **maroon**-filled cells represent the brain areas that were predictive of SCR-based learning outcome across 4 imaging modalities; **yellow**-filled cells represent the brain areas that were predictive of SCR-based learning outcome across 3 imaging modalities; **cyan**-filled cells represent the brain areas that were predictive of SCR-based learning outcome across 2 imaging modalities; **blue**-filled cells represent the brain areas that were predictive of SCR-based learning outcome in 1 imaging modality. The 'L_' and 'R_' prefixes to region codes correspond to the left hemisphere and the right hemisphere. The table with longer anatomical names of the corresponding coded areas is presented at the end of the supplementary materials, Table S7.

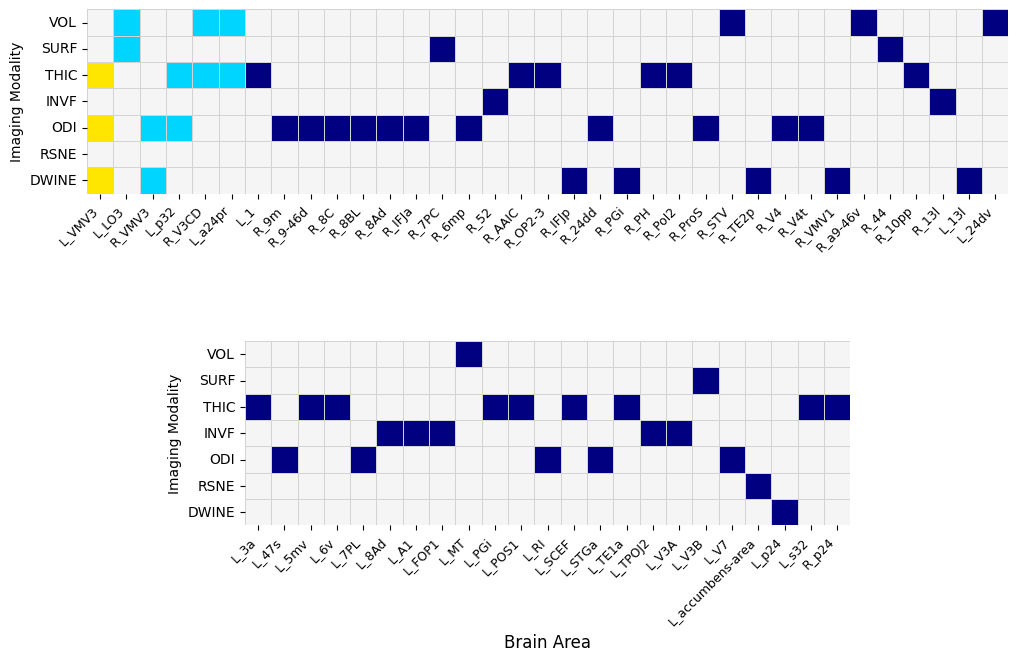

**Figure S4.** A schematic, visual depiction of the brain areas in each imaging modality that were consistently predictive of the SCR outcome during extinction learning across at least 50% of 500 bootstrap iterations of cross-validated second-level, multimodal ElasticNet, which received the shortlisted brain areas from all imaging modalities of an earlier 1000 iteration bootstrapped cross-validation in Figure S2. The **maroon**-filled cells represent the brain areas that were predictive of SCR-based learning outcome across 4 imaging modalities; **yellow**-filled cells represent the brain areas that were predictive of SCR-based learning outcome across 3 imaging modalities; **cyan**-filled cells represent the brain areas that were predictive of SCR-based learning outcome across 2 imaging modalities; **blue**-filled cells represent the brain areas that were predictive of SCR-based learning outcome in 1 imaging modality. The 'L_' and 'R_' prefixes to region codes correspond to the left hemisphere and the right hemisphere. The table with longer anatomical names of the corresponding coded areas is presented at the end of the supplementary materials, Table S7.

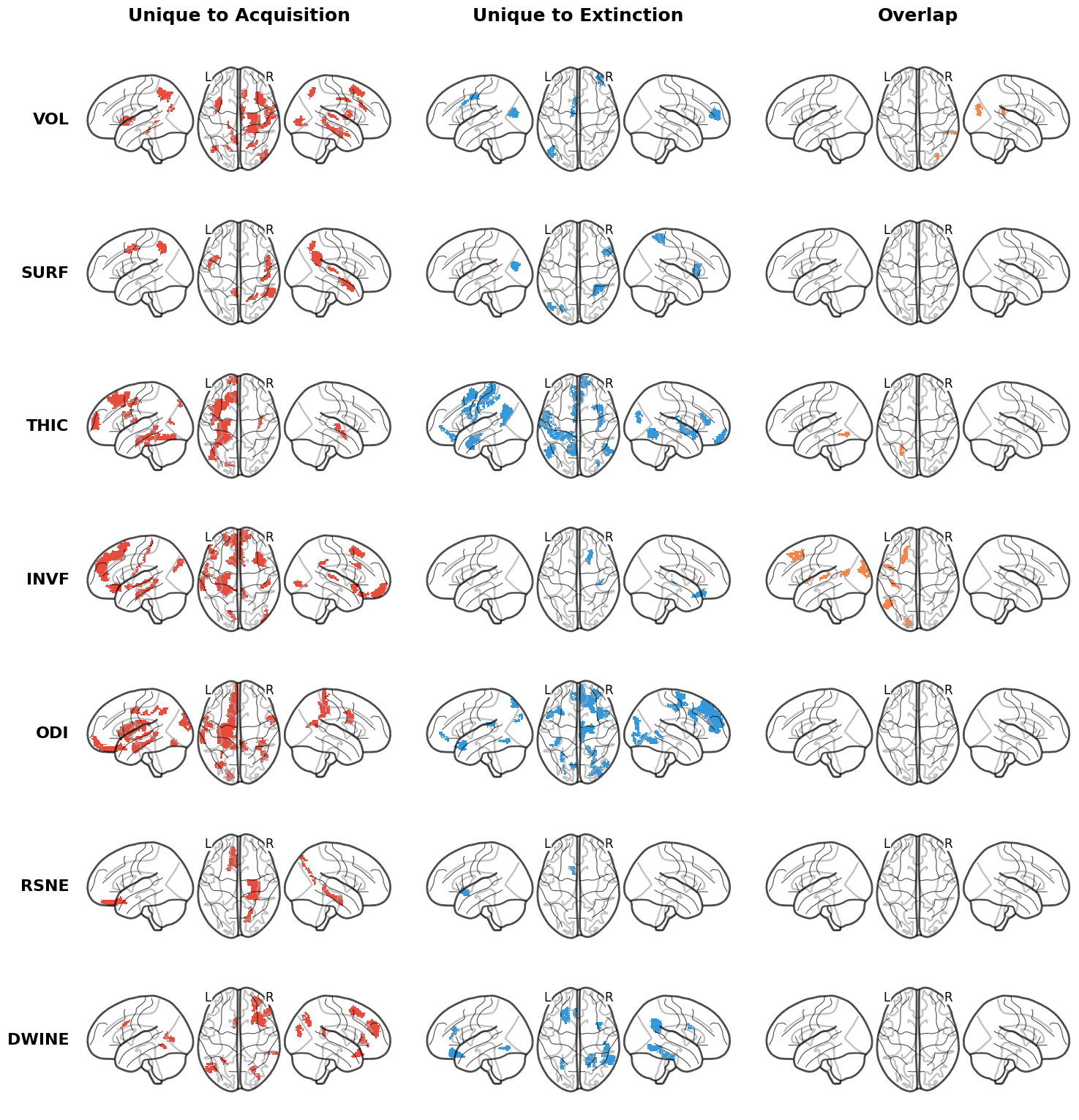

**Figure S5. Modality-specific brain regions predictive of physiological fear responses across seven neuroimaging modalities from a more conservative, second-level multimodal ElasticNet regression.**

This figure mirrors Figure 3 but is based on a second-level (stacked) ElasticNet model rather than independent per-modality fits. The second-level model received as input only those brain regions that survived the stability threshold (≥50% of 1,000 bootstrap iterations) in the first-level analysis (Supplementary Figures S1–S2), and was itself bootstrapped across 500 iterations with nested cross-validation. This two-stage filtering imposes a more stringent selection criterion, retaining only regions whose predictive contribution is stable both within and across modelling stages. Each row corresponds to one imaging modality (VOL = regional brain volume; SURF = cortical surface area; THIC = cortical thickness; INVF = intra-neurite volume fraction; ODI = orientation dispersion index; RSNE = resting-state connectivity nodal efficiency; DWINE = diffusion-weighted imaging connectivity nodal efficiency). Columns display regions consistently predictive of differential SCRs during fear acquisition (left, red), regions consistently predictive of the difference of the differential SCRs over the course of fear extinction (middle, blue), and the overlapping regions that were predictive during both phases (right, orange). Glass brain views show left lateral, dorsal, and right lateral perspectives for each condition. The complete list of brain areas and their bootstrap stability statistics is provided in Supplementary Tables S1 and S2, and visualised as heatmaps in Supplementary Figures S3 and S4.

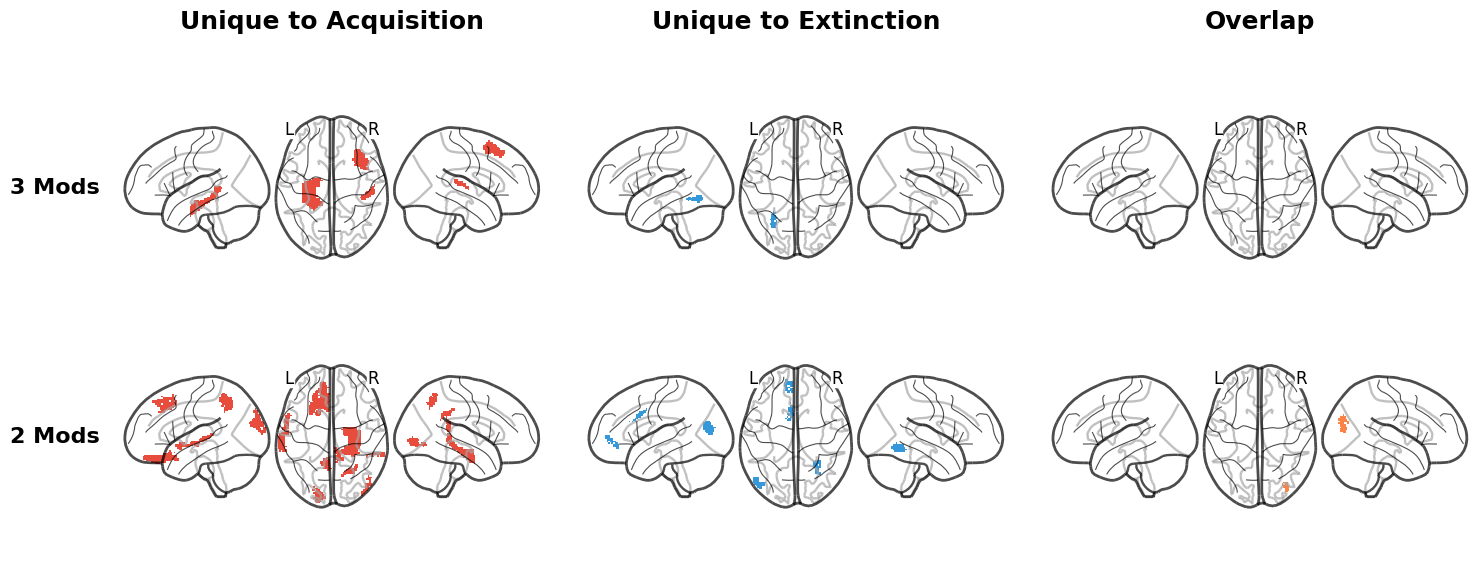

**Figure S6. Brain regions whose predictive relationship with fear learning replicates across multiple neuroimaging modalities, from a more conservative, second-level multimodal ElasticNet regression.**

This figure mirrors Figure 4 but is derived from the second-level stacked model (500 bootstrap iterations), which operated on the pre-filtered set of regions from the first-level analysis (Supplementary Figures S1–S2). Cross-modal consensus was assessed identically to Figure 4: rows indicate regions identified as stable predictors in at least three (top) or two (bottom) distinct imaging modalities. No region reached the four-modality threshold in the second-level model. Because the input features were already pre-screened for stability at the first level, consensus at this stage reflects a doubly conservative criterion: regions must have been stable in the first-level per-modality fits and again contribute reliably in the pooled second-level multimodal model. Color coding and column arrangement follow the convention of Figure 4 (red = fear acquisition; blue = extinction; orange = dynamic/overlap). The complete list of brain areas and their corresponding consensus levels is provided in Supplementary Figures S3 and S4, and Supplementary Tables S1 and S2.

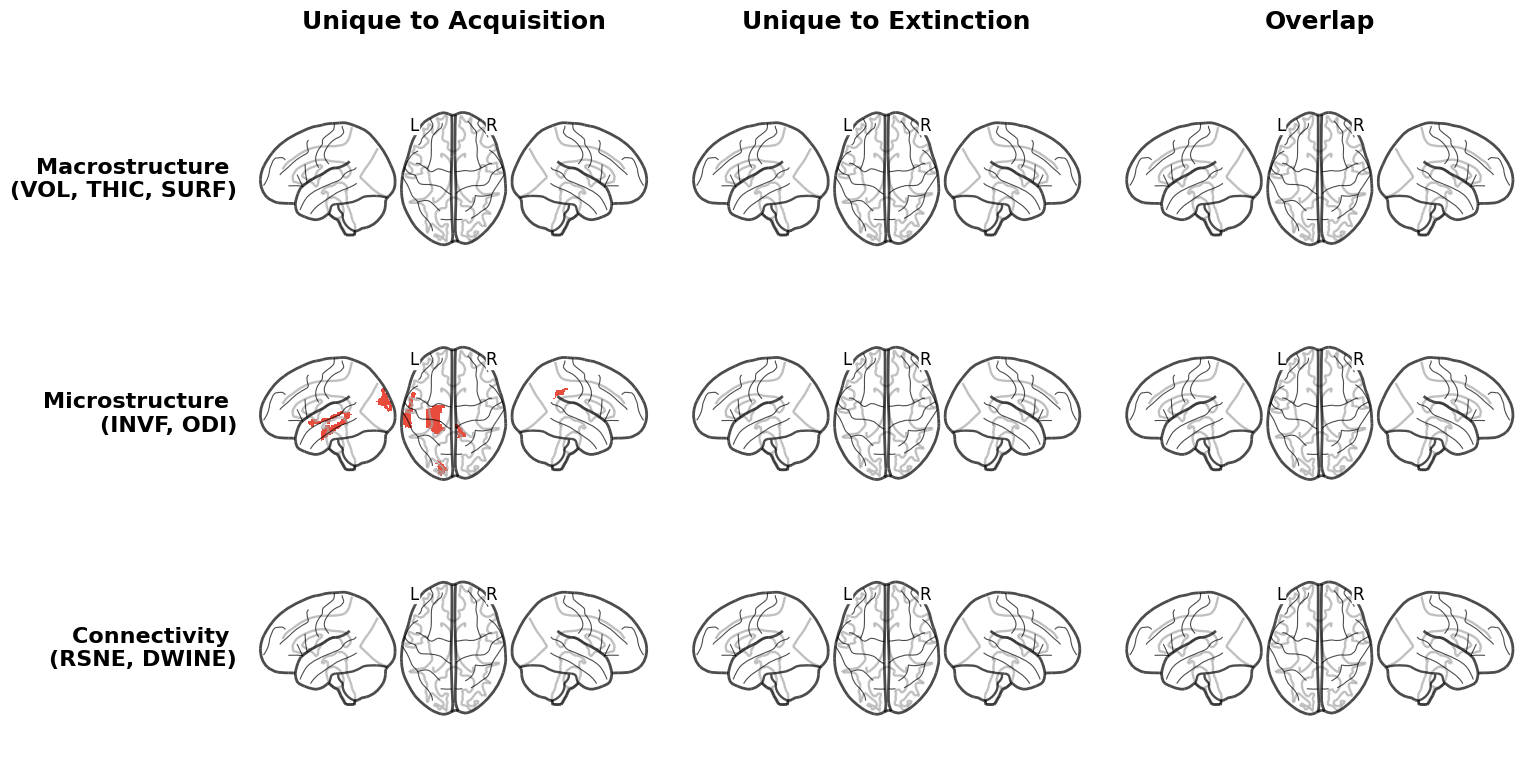

**Figure S7. Convergence of fear learning predictors grouped across three neuroimaging domains: macrostructure, microstructure, and connectivity, from a more conservative, second-level multimodal ElasticNet regression.**

This figure mirrors Figure 5 but is derived from the second-level stacked model (500 bootstrap iterations), using the pre-filtered region set from the first-level analysis (Supplementary Figures S1–S2) as input. As in Figure 5, modalities were grouped into three domains: macrostructure (VOL, SURF, THIC), microstructure (INVF, ODI), and connectivity (RSNE, DWINE). The region is shown if it received stable non-zero weights (≥50% of 500 bootstrap iterations) by all modalities, indicating convergent evidence that is unlikely to reflect modality-specific measurement variance. This two-stage design means that regions shown here have converging evidence from both within-modality and cross-modality analyses, making them the most robustly supported predictors reported in this study. Rows represent specific domain-pair and domain-triple overlaps; columns and colour coding follow Figures 3–5 (red = fear acquisition; blue = extinction; orange = dynamic). Glass brain views show left lateral, dorsal, and right lateral perspectives. The full list of brain areas within each domain grouping is provided in Supplementary Figures S3 and S4.

**Table S1**. Stability of the brain areas across 1000 bootstrap iterations for fear acquisition training data of a first-level ElasticNet regression. The **Modality** column represents the imaging modality from which the data have been used to fit the ElasticNet regression. The **Brain Area** column corresponds to brain areas of cortical and, where applicable, subcortical brain areas according to HCPMMP and FreeSurfer's subcortical automated labeling. The 'L_' and 'R_' prefixes to region codes correspond to the left hemisphere and the right hemisphere. The table with longer anatomical names of the corresponding coded areas is presented at the end of the supplementary materials, Table S7. The **Stability %** shows the percentage of bootstrap iterations where this specific brain area has received a non-zero weight as a result of nested 5-fold cross-validation in predicting the differential (CS+ - CS–) SCR outcome of fear learning. The **Mean Coefficient**, **Sign Consistency,** and **Sign Directionality** show the average beta coefficient of the regularized ElasticNet Regression, consistency of sign, and the directionality (positive or negative) of the beta weight across 5-fold cross-validation in each of 1000 bootstrap folds. The rows are sorted by the **Stability %** column in decreasing order for each of the imaging modalities. The list of brain areas is thresholded with at least 50% stability across 1000 bootstrap iterations (or being consistently nonzero in at least 500 iterations).

| **Modality** | **Brain Area** | **Stability %** | **Mean Coefficient** | **Sign Consistency** | **Sign Directionality** |
| --- | --- | --- | --- | --- | --- |
| VOL | R_PreS | 0.820 | -0.06501 | 1.000 | − |
|  | R_p24pr | 0.770 | 0.06321 | 1.000 | + |
|  | R_H | 0.735 | -0.05701 | 1.000 | − |
|  | L_TPOJ3 | 0.712 | -0.04476 | 1.000 | − |
|  | R_V8 | 0.712 | 0.04752 | 1.000 | + |
|  | L_PCV | 0.711 | 0.04751 | 1.000 | + |
|  | L_MIP | 0.705 | -0.05024 | 1.000 | − |
|  | L_IFJa | 0.701 | 0.05361 | 0.990 | + |
|  | R_MIP | 0.698 | 0.04592 | 0.999 | + |
|  | R_MI | 0.637 | -0.06301 | 1.000 | − |
|  | R_8Av | 0.635 | -0.08092 | 0.969 | − |
|  | R_V3CD | 0.628 | 0.06909 | 0.995 | + |
|  | L_MI | 0.602 | -0.06070 | 1.000 | − |
|  | L_PreS | 0.598 | -0.04946 | 1.000 | − |
|  | R_6v | 0.567 | -0.05383 | 0.996 | − |
|  | R_LO1 | 0.562 | 0.05228 | 0.998 | + |
|  | R_LBelt | 0.560 | 0.06628 | 0.996 | + |
|  | R_STSva | 0.560 | -0.05742 | 0.998 | − |
|  | R_STV | 0.559 | 0.06273 | 0.959 | + |
|  | R_TA2 | 0.542 | -0.04164 | 1.000 | − |
|  | R_7PL | 0.532 | 0.05867 | 0.930 | + |
|  | R_LO2 | 0.525 | -0.03912 | 1.000 | − |
|  | L_H | 0.520 | -0.07003 | 0.973 | − |
|  | R_43 | 0.520 | -0.06156 | 0.985 | − |
|  | R_a32pr | 0.520 | 0.04003 | 0.967 | + |
|  | R_PeEc | 0.517 | -0.04200 | 0.911 | − |
| SURF | R_LBelt | 0.800 | 0.08439 | 1.000 | + |
|  | R_PreS | 0.744 | -0.06754 | 1.000 | − |
|  | R_MIP | 0.732 | 0.06725 | 0.995 | + |
|  | R_PeEc | 0.687 | -0.07232 | 0.997 | − |
|  | L_MST | 0.650 | -0.08396 | 0.989 | − |
|  | R_H | 0.627 | -0.05260 | 1.000 | − |
|  | R_PGi | 0.620 | -0.05483 | 0.995 | − |
|  | L_PCV | 0.617 | 0.05787 | 1.000 | + |
|  | R_44 | 0.615 | 0.06024 | 1.000 | + |
|  | R_STSda | 0.580 | -0.08186 | 0.993 | − |
|  | R_LIPd | 0.560 | 0.04959 | 1.000 | + |
|  | R_8Av | 0.554 | -0.06173 | 0.986 | − |
|  | L_55b | 0.542 | 0.07172 | 0.967 | + |
|  | R_6v | 0.542 | -0.05304 | 0.996 | − |
|  | R_p24pr | 0.535 | 0.04541 | 1.000 | + |
|  | R_FOP1 | 0.513 | 0.05299 | 0.998 | + |
|  | L_ProS | 0.505 | 0.05714 | 0.988 | + |
|  | R_PCV | 0.502 | 0.05611 | 0.998 | + |
| THIC | R_PoI1 | 0.818 | 0.08352 | 1.000 | + |
|  | L_a24 | 0.753 | -0.06991 | 0.999 | − |
|  | L_FOP2 | 0.716 | -0.07400 | 0.999 | − |
|  | L_V6A | 0.655 | 0.04565 | 1.000 | + |
|  | L_10d | 0.640 | 0.06948 | 1.000 | + |
|  | L_VMV3 | 0.638 | 0.09927 | 0.987 | + |
|  | L_PHA3 | 0.633 | -0.05208 | 0.995 | − |
|  | L_MT | 0.625 | 0.05979 | 0.995 | + |
|  | L_H | 0.588 | -0.06955 | 0.974 | − |
|  | L_MI | 0.581 | -0.06518 | 0.991 | − |
|  | L_8BL | 0.578 | -0.07583 | 0.998 | − |
|  | R_V3CD | 0.574 | 0.04796 | 0.997 | + |
|  | L_FEF | 0.562 | 0.07927 | 0.996 | + |
|  | L_8Av | 0.561 | -0.05399 | 1.000 | − |
|  | L_V3A | 0.549 | 0.04254 | 0.998 | + |
|  | R_7m | 0.546 | -0.04366 | 0.998 | − |
|  | L_9a | 0.540 | -0.05866 | 0.996 | − |
|  | L_FFC | 0.522 | 0.04010 | 1.000 | + |
|  | L_8Ad | 0.514 | 0.06401 | 0.994 | + |
|  | L_IFJp | 0.510 | -0.03545 | 1.000 | − |
|  | L_IFJa | 0.502 | 0.03826 | 0.994 | + |
| INVF | L_A5 | 0.794 | -0.06515 | 1.000 | − |
|  | R_a24pr | 0.776 | 0.04662 | 1.000 | + |
|  | R_8Av | 0.762 | 0.05479 | 1.000 | + |
|  | R_OP1 | 0.754 | 0.05506 | 1.000 | + |
|  | R_PCV | 0.727 | -0.06034 | 1.000 | − |
|  | L_8BM | 0.723 | 0.07435 | 1.000 | + |
|  | L_TPOJ2 | 0.709 | 0.08386 | 1.000 | + |
|  | L_8Ad | 0.708 | -0.08480 | 0.999 | − |
|  | R_10v | 0.707 | -0.04456 | 1.000 | − |
|  | L_V6 | 0.698 | -0.04566 | 1.000 | − |
|  | L_FOP1 | 0.689 | -0.08861 | 0.997 | − |
|  | L_V6A | 0.681 | 0.05465 | 0.997 | + |
|  | R_31pv | 0.678 | -0.05709 | 1.000 | − |
|  | L_V3A | 0.671 | 0.06700 | 0.996 | + |
|  | R_A1 | 0.670 | 0.03613 | 0.775 | + |
|  | R_6ma | 0.667 | 0.03976 | 1.000 | + |
|  | L_H | 0.655 | -0.05198 | 0.994 | − |
|  | L_3a | 0.650 | 0.06728 | 0.988 | + |
|  | R_VMV2 | 0.649 | -0.03865 | 0.995 | − |
|  | L_V3B | 0.646 | -0.05955 | 1.000 | − |
|  | R_8BL | 0.635 | 0.04114 | 1.000 | + |
|  | L_2 | 0.635 | 0.06812 | 0.994 | + |
|  | R_47s | 0.634 | 0.04125 | 0.998 | + |
|  | L_A1 | 0.609 | -0.07292 | 0.962 | − |
|  | R_10pp | 0.602 | -0.03518 | 1.000 | − |
|  | R_STSdp | 0.571 | -0.03636 | 1.000 | − |
|  | L_46 | 0.569 | -0.05291 | 1.000 | − |
|  | L_9a | 0.556 | -0.04959 | 0.998 | − |
|  | R_VVC | 0.551 | -0.06001 | 0.993 | − |
|  | L_47m | 0.550 | 0.04785 | 0.987 | + |
|  | L_SFL | 0.540 | 0.03901 | 1.000 | + |
|  | R_9a | 0.539 | 0.03297 | 0.998 | + |
|  | L_47l | 0.534 | 0.05488 | 0.989 | + |
|  | R_AAIC | 0.532 | 0.04443 | 0.893 | + |
|  | L_9m | 0.531 | 0.04506 | 0.996 | + |
|  | L_10r | 0.508 | 0.04481 | 0.982 | + |
|  | L_VIP | 0.505 | -0.04979 | 0.958 | − |
|  | R_LBelt | 0.504 | 0.03028 | 0.954 | + |
|  | R_5mv | 0.504 | 0.04724 | 0.984 | + |
|  | R_LO2 | 0.502 | -0.03675 | 0.970 | − |
| ODI | L_V3A | 0.831 | -0.07238 | 0.998 | − |
|  | R_6v | 0.767 | 0.06553 | 0.999 | + |
|  | R_5m | 0.734 | -0.04384 | 1.000 | − |
|  | L_A5 | 0.733 | -0.06829 | 1.000 | − |
|  | L_H | 0.665 | -0.04224 | 0.998 | − |
|  | L_LIPd | 0.662 | 0.06329 | 0.998 | + |
|  | L_52 | 0.648 | -0.03995 | 0.998 | − |
|  | L_IP2 | 0.647 | 0.04983 | 0.997 | + |
|  | L_OFC | 0.644 | 0.05010 | 1.000 | + |
|  | L_caudate | 0.626 | 0.04105 | 0.995 | + |
|  | R_TPOJ3 | 0.626 | -0.05876 | 0.997 | − |
|  | L_43 | 0.611 | -0.05022 | 1.000 | − |
|  | R_31pv | 0.576 | 0.08164 | 0.997 | + |
|  | R_PreS | 0.564 | -0.04295 | 0.991 | − |
|  | R_EC | 0.550 | -0.06139 | 0.991 | − |
|  | R_Ig | 0.549 | -0.04973 | 0.996 | − |
|  | L_V8 | 0.548 | 0.04539 | 1.000 | + |
|  | R_LIPv | 0.547 | 0.03912 | 0.993 | + |
|  | L_Ig | 0.531 | 0.02964 | 0.699 | + |
|  | L_10v | 0.515 | -0.03398 | 0.998 | − |
|  | R_AIP | 0.510 | 0.03007 | 1.000 | + |
|  | L_24dv | 0.510 | -0.06562 | 0.967 | − |
|  | L_amygdala | 0.505 | 0.03004 | 1.000 | + |
| RSNE | R_s32 | 0.692 | 0.09996 | 0.986 | + |
|  | R_H | 0.668 | -0.06342 | 0.981 | − |
|  | L_TGv | 0.582 | -0.16127 | 0.790 | − |
|  | L_accumbens-area | 0.580 | 0.10253 | 0.859 | + |
|  | R_SFL | 0.572 | -0.11864 | 0.930 | − |
|  | L_OFC | 0.564 | -0.07080 | 0.995 | − |
|  | L_TPOJ2 | 0.554 | -0.10544 | 1.000 | − |
|  | R_45 | 0.552 | 0.11264 | 1.000 | + |
|  | L_amygdala | 0.549 | -0.08709 | 0.996 | − |
|  | R_DVT | 0.545 | 0.14685 | 1.000 | + |
|  | L_PeEc | 0.521 | 0.10787 | 0.994 | + |
|  | L_8BL | 0.503 | 0.07932 | 0.988 | + |
|  | R_AAIC | 0.502 | -0.15064 | 0.982 | − |
| DWINE | R_FOP1 | 0.783 | 0.08021 | 1.000 | + |
|  | R_V3CD | 0.761 | 0.09773 | 1.000 | + |
|  | R_LO1 | 0.706 | 0.06673 | 0.994 | + |
|  | L_i6-8 | 0.705 | 0.05489 | 1.000 | + |
|  | R_IP0 | 0.677 | -0.07225 | 1.000 | − |
|  | L_VMV2 | 0.644 | 0.05481 | 0.997 | + |
|  | R_V6 | 0.642 | -0.05712 | 1.000 | − |
|  | R_45 | 0.629 | 0.07237 | 1.000 | + |
|  | L_MI | 0.623 | -0.04914 | 1.000 | − |
|  | R_IPS1 | 0.616 | -0.03860 | 1.000 | − |
|  | R_PGi | 0.577 | -0.03960 | 0.979 | − |
|  | R_STV | 0.574 | 0.05305 | 0.911 | + |
|  | L_FST | 0.566 | -0.05503 | 1.000 | − |
|  | R_8Av | 0.566 | -0.06818 | 0.890 | − |
|  | R_9-46d | 0.557 | -0.04022 | 1.000 | − |
|  | L_a24pr | 0.556 | 0.06429 | 0.894 | + |
|  | R_47s | 0.552 | -0.04545 | 0.998 | − |
|  | L_8Av | 0.547 | 0.05233 | 0.996 | + |
|  | R_POS2 | 0.521 | -0.04439 | 1.000 | − |
|  | R_FOP3 | 0.520 | 0.05122 | 0.994 | + |
|  | L_7Am | 0.520 | 0.04761 | 1.000 | + |
|  | R_p32 | 0.518 | -0.03622 | 1.000 | − |
|  | L_LIPd | 0.515 | -0.04544 | 0.990 | − |
|  | R_3a | 0.512 | -0.03377 | 1.000 | − |
|  | R_PFop | 0.512 | -0.04622 | 1.000 | − |
|  | R_s32 | 0.511 | -0.02735 | 1.000 | − |
|  | R_LBelt | 0.511 | 0.04521 | 0.998 | + |
|  | R_TE2a | 0.509 | 0.05019 | 0.996 | + |
|  | R_EC | 0.507 | -0.05492 | 0.996 | − |
|  | L_MST | 0.503 | -0.03859 | 0.998 | − |

**Table S2**. Stability of the brain areas across 1000 bootstrap iterations for extinction training data of a first-level ElasticNet regression. The **Modality** column represents the imaging modality from which the data have been used to fit the ElasticNet regression. The **Brain Area** column corresponds to brain areas of cortical, and where applicable, subcortical brain areas according to HCPMMP and FreeSurfer's subcortical automated labeling. The 'L_' and 'R_' prefixes to region codes correspond to the left hemisphere and the right hemisphere. The table with longer anatomical names of the corresponding coded areas is presented at the end of the supplementary materials, Table S7. The **Stability %** shows the percentage of bootstrap iterations where this specific brain area has received a non-zero weight as a result of nested 5-fold cross-validation in predicting a slope of a difference ((Early CS+ - Early CS–) - (Late CS+ - Late CS–)) SCR outcome during extinction learning. The **Mean Coefficient**, **Sign Consistency,** and **Sign Directionality** show the average beta coefficient of the regularized ElasticNet Regression, consistency of sign, and the directionality (positive or negative) of the beta weight across 5-fold cross-validation in each of 1000 bootstrap folds. The rows are sorted by the **Stability %** column in decreasing order for each of the imaging modalities. The list of brain areas is thresholded with at least 50% stability across 1000 bootstrap iterations (or being consistently nonzero in at least 500 iterations).

| **Modality** | **Brain Area** | **Stability %** | **Mean Coefficient** | **Sign Consistency** | **Sign Directionality** |
| --- | --- | --- | --- | --- | --- |
| VOL | L_V3B | 0.820 | -0.07056 | 1.000 | − |
|  | L_a24pr | 0.805 | 0.08011 | 0.999 | + |
|  | R_a9-46v | 0.794 | 0.07232 | 0.999 | + |
|  | R_8Av | 0.780 | -0.09037 | 1.000 | − |
|  | L_MT | 0.736 | 0.05687 | 1.000 | + |
|  | R_STV | 0.732 | 0.07913 | 0.990 | + |
|  | R_9p | 0.730 | 0.05907 | 1.000 | + |
|  | L_p32 | 0.716 | 0.06807 | 0.996 | + |
|  | L_24dv | 0.701 | 0.06168 | 1.000 | + |
|  | L_10r | 0.673 | -0.06335 | 0.999 | − |
|  | L_LO3 | 0.672 | 0.07008 | 1.000 | + |
|  | L_a10p | 0.660 | 0.06175 | 0.994 | + |
|  | L_PF | 0.657 | -0.07610 | 1.000 | − |
|  | L_STSvp | 0.654 | -0.05985 | 0.998 | − |
|  | R_47s | 0.603 | -0.04879 | 1.000 | − |
|  | R_6a | 0.554 | -0.05089 | 0.998 | − |
|  | R_H | 0.545 | -0.04063 | 1.000 | − |
|  | L_EC | 0.542 | 0.05660 | 1.000 | + |
|  | R_7Pm | 0.540 | 0.05529 | 0.994 | + |
|  | R_V3CD | 0.538 | 0.07230 | 1.000 | + |
|  | R_24dv | 0.536 | -0.04248 | 0.993 | − |
|  | L_a9-46v | 0.513 | -0.05547 | 1.000 | − |
|  | L_31a | 0.511 | 0.05344 | 0.998 | + |
|  | R_MI | 0.504 | -0.05177 | 0.996 | − |
| SURF | R_PoI1 | 0.767 | -0.09807 | 1.000 | − |
|  | L_LO3 | 0.754 | 0.08108 | 1.000 | + |
|  | R_9p | 0.742 | 0.06685 | 1.000 | + |
|  | L_TPOJ1 | 0.722 | 0.07607 | 1.000 | + |
|  | L_55b | 0.700 | 0.08135 | 0.994 | + |
|  | L_V3B | 0.695 | -0.06169 | 1.000 | − |
|  | L_STSvp | 0.669 | -0.06463 | 0.999 | − |
|  | L_10v | 0.666 | -0.06675 | 1.000 | − |
|  | R_44 | 0.662 | 0.06728 | 1.000 | + |
|  | L_31a | 0.653 | 0.09094 | 1.000 | + |
|  | R_IFJa | 0.646 | -0.06356 | 1.000 | − |
|  | R_TGd | 0.643 | 0.05506 | 1.000 | + |
|  | R_a10p | 0.628 | 0.06001 | 1.000 | + |
|  | R_8Av | 0.615 | -0.07248 | 1.000 | − |
|  | L_PSL | 0.605 | -0.05793 | 0.992 | − |
|  | R_6a | 0.572 | -0.05913 | 1.000 | − |
|  | L_8C | 0.559 | 0.06270 | 1.000 | + |
|  | L_FOP1 | 0.547 | 0.05653 | 0.978 | + |
|  | L_VMV2 | 0.538 | -0.05164 | 0.994 | − |
|  | L_43 | 0.531 | 0.06452 | 0.998 | + |
|  | L_DVT | 0.528 | -0.06364 | 0.985 | − |
|  | R_a9-46v | 0.527 | 0.05444 | 0.998 | + |
|  | R_7PC | 0.523 | -0.04578 | 0.994 | − |
|  | R_FEF | 0.521 | -0.04326 | 0.994 | − |
|  | R_MIP | 0.520 | 0.04854 | 0.996 | + |
|  | L_IFJp | 0.518 | -0.06963 | 0.998 | − |
|  | L_OP4 | 0.518 | 0.04859 | 1.000 | + |
|  | R_STSda | 0.506 | -0.06207 | 0.990 | − |
| THIC | L_5mv | 0.923 | -0.11756 | 1.000 | − |
|  | R_PoI2 | 0.858 | 0.05344 | 1.000 | + |
|  | L_VMV3 | 0.793 | 0.08680 | 0.999 | + |
|  | L_p32 | 0.756 | 0.06876 | 1.000 | + |
|  | L_MT | 0.754 | 0.06300 | 0.999 | + |
|  | L_TE1a | 0.752 | -0.06822 | 1.000 | − |
|  | R_p24 | 0.709 | -0.05007 | 1.000 | − |
|  | L_SCEF | 0.708 | 0.05172 | 1.000 | + |
|  | L_3a | 0.688 | -0.06751 | 1.000 | − |
|  | L_PGi | 0.687 | 0.05551 | 1.000 | + |
|  | L_a24pr | 0.686 | 0.05513 | 1.000 | + |
|  | R_AAIC | 0.648 | 0.04695 | 1.000 | + |
|  | L_s32 | 0.631 | -0.04589 | 0.998 | − |
|  | R_OP2-3 | 0.606 | 0.04639 | 0.997 | + |
|  | L_V7 | 0.595 | 0.04794 | 1.000 | + |
|  | R_STV | 0.590 | 0.04601 | 1.000 | + |
|  | R_V3CD | 0.585 | 0.04378 | 0.998 | + |
|  | R_8C | 0.577 | 0.04104 | 1.000 | + |
|  | L_POS1 | 0.571 | -0.05243 | 0.998 | − |
|  | L_6v | 0.568 | -0.05664 | 0.998 | − |
|  | L_1 | 0.551 | 0.05498 | 1.000 | + |
|  | R_10pp | 0.544 | -0.03585 | 1.000 | − |
|  | R_PH | 0.530 | -0.03996 | 0.989 | − |
|  | L_PH | 0.515 | -0.04221 | 0.996 | − |
| INVF | R_13l | 0.815 | 0.08012 | 1.000 | + |
|  | L_PoI2 | 0.799 | -0.06202 | 1.000 | − |
|  | L_TPOJ2 | 0.758 | 0.09569 | 1.000 | + |
|  | R_52 | 0.739 | 0.06547 | 0.999 | + |
|  | L_A1 | 0.734 | -0.09765 | 1.000 | − |
|  | R_OP1 | 0.721 | 0.06339 | 0.999 | + |
|  | L_FOP1 | 0.720 | -0.08200 | 1.000 | − |
|  | L_PHA1 | 0.712 | -0.06242 | 1.000 | − |
|  | L_PI | 0.710 | -0.05515 | 0.994 | − |
|  | L_52 | 0.708 | -0.06702 | 1.000 | − |
|  | L_V3B | 0.655 | -0.06904 | 0.998 | − |
|  | L_TE2a | 0.655 | 0.06734 | 1.000 | + |
|  | R_TE2p | 0.650 | 0.05157 | 1.000 | + |
|  | L_47m | 0.644 | 0.07406 | 0.981 | + |
|  | L_V3A | 0.612 | 0.04744 | 0.998 | + |
|  | L_TE2p | 0.570 | 0.05154 | 0.996 | + |
|  | L_4 | 0.564 | 0.08848 | 0.993 | + |
|  | R_9m | 0.556 | -0.04597 | 1.000 | − |
|  | R_TGv | 0.553 | 0.04094 | 0.987 | + |
|  | L_TPOJ3 | 0.539 | 0.03761 | 1.000 | + |
|  | L_POS1 | 0.526 | 0.05122 | 1.000 | + |
|  | L_7PC | 0.523 | 0.05626 | 1.000 | + |
|  | L_5m | 0.519 | -0.05058 | 0.927 | − |
|  | R_PI | 0.516 | -0.04174 | 0.986 | − |
|  | L_OP2-3 | 0.511 | 0.04963 | 0.998 | + |
|  | L_7AL | 0.508 | -0.03752 | 0.998 | − |
|  | R_IP0 | 0.505 | -0.03958 | 1.000 | − |
| ODI | L_47s | 0.844 | 0.08310 | 1.000 | + |
|  | L_p32 | 0.836 | -0.07649 | 1.000 | − |
|  | R_9-46d | 0.819 | 0.06719 | 1.000 | + |
|  | L_a10p | 0.778 | -0.05366 | 1.000 | − |
|  | R_ProS | 0.752 | -0.06046 | 1.000 | − |
|  | R_8C | 0.728 | -0.05756 | 1.000 | − |
|  | R_EC | 0.700 | -0.06557 | 0.999 | − |
|  | L_V7 | 0.682 | -0.05936 | 1.000 | − |
|  | R_9m | 0.675 | 0.05063 | 1.000 | + |
|  | L_VMV3 | 0.669 | -0.06260 | 0.996 | − |
|  | L_STGa | 0.660 | -0.06455 | 1.000 | − |
|  | R_V4t | 0.657 | 0.04389 | 1.000 | + |
|  | R_8Ad | 0.653 | -0.04381 | 1.000 | − |
|  | R_24dd | 0.600 | 0.03951 | 0.998 | + |
|  | R_6mp | 0.566 | 0.04200 | 1.000 | + |
|  | R_IFJa | 0.565 | -0.04417 | 0.998 | − |
|  | L_PHA2 | 0.565 | -0.04110 | 1.000 | − |
|  | L_7PL | 0.562 | -0.05008 | 1.000 | − |
|  | R_V4 | 0.539 | -0.03055 | 1.000 | − |
|  | R_52 | 0.528 | -0.03850 | 0.996 | − |
|  | R_VMV3 | 0.524 | -0.03293 | 1.000 | − |
|  | L_PF | 0.523 | 0.06217 | 0.996 | + |
|  | L_a24pr | 0.516 | -0.04108 | 1.000 | − |
|  | L_RI | 0.511 | 0.04349 | 0.986 | + |
|  | R_6a | 0.503 | -0.04031 | 0.998 | − |
| RSNE | L_accumbens-area | 0.940 | 0.17295 | 1.000 | + |
|  | L_IP0 | 0.924 | 0.20337 | 1.000 | + |
|  | L_i6-8 | 0.886 | -0.18307 | 1.000 | − |
|  | R_IFSp | 0.873 | 0.15094 | 1.000 | + |
|  | L_H | 0.818 | -0.07148 | 0.989 | − |
|  | L_10pp | 0.772 | -0.09825 | 0.997 | − |
|  | R_PreS | 0.748 | -0.12809 | 0.999 | − |
|  | R_33pr | 0.746 | 0.09529 | 0.996 | + |
|  | L_Pir | 0.656 | -0.08897 | 1.000 | − |
|  | R_45 | 0.591 | 0.08919 | 0.998 | + |
|  | L_TGv | 0.578 | -0.05993 | 0.825 | − |
|  | L_IP1 | 0.576 | -0.13848 | 0.997 | − |
|  | R_31pd | 0.576 | 0.08978 | 1.000 | + |
|  | R_5L | 0.567 | 0.08926 | 0.979 | + |
|  | L_TPOJ3 | 0.551 | -0.08215 | 0.980 | − |
|  | R_pallidum | 0.540 | -0.07205 | 0.931 | − |
|  | R_PGp | 0.525 | -0.08160 | 1.000 | − |
|  | R_5mv | 0.524 | 0.10228 | 1.000 | + |
|  | L_PreS | 0.521 | -0.05967 | 0.990 | − |
|  | R_p24 | 0.519 | 0.07005 | 1.000 | + |
|  | R_AAIC | 0.518 | -0.09366 | 0.981 | − |
|  | R_7AL | 0.514 | 0.06921 | 0.998 | + |
|  | R_s32 | 0.508 | 0.04613 | 0.878 | + |
|  | R_5m | 0.506 | 0.08731 | 0.998 | + |
| DWINE | R_IPS1 | 0.785 | -0.05744 | 1.000 | − |
|  | L_TA2 | 0.773 | 0.07170 | 0.999 | + |
|  | R_PSL | 0.769 | 0.05986 | 1.000 | + |
|  | L_FFC | 0.758 | 0.08555 | 0.997 | + |
|  | R_IFSp | 0.745 | -0.05440 | 1.000 | − |
|  | L_IP1 | 0.738 | 0.06347 | 1.000 | + |
|  | L_TE1a | 0.719 | 0.05267 | 0.999 | + |
|  | L_PF | 0.687 | -0.07277 | 1.000 | − |
|  | R_PHA3 | 0.680 | -0.04899 | 1.000 | − |
|  | L_6mp | 0.679 | 0.07590 | 0.999 | + |
|  | R_9-46d | 0.674 | -0.04741 | 1.000 | − |
|  | R_8Av | 0.657 | -0.08255 | 1.000 | − |
|  | L_p24 | 0.636 | 0.05618 | 0.992 | + |
|  | R_VMV1 | 0.635 | -0.04879 | 1.000 | − |
|  | R_STGa | 0.616 | 0.04628 | 1.000 | + |
|  | R_TE2p | 0.597 | -0.04432 | 0.993 | − |
|  | R_PGi | 0.595 | -0.04442 | 0.982 | − |
|  | L_13l | 0.587 | -0.04877 | 1.000 | − |
|  | R_V3CD | 0.578 | 0.05003 | 0.995 | + |
|  | L_a47r | 0.574 | -0.05771 | 1.000 | − |
|  | L_i6-8 | 0.568 | 0.04872 | 0.991 | + |
|  | L_PoI2 | 0.567 | 0.04825 | 1.000 | + |
|  | L_STV | 0.566 | 0.05349 | 0.989 | + |
|  | L_VMV3 | 0.555 | -0.05298 | 0.995 | − |
|  | R_VMV3 | 0.555 | -0.04660 | 0.996 | − |
|  | R_s6-8 | 0.523 | 0.03788 | 0.998 | + |
|  | L_PFop | 0.519 | -0.04378 | 1.000 | − |
|  | L_6r | 0.511 | 0.05676 | 1.000 | + |
|  | R_V4t | 0.507 | -0.03340 | 1.000 | − |
|  | R_1 | 0.505 | -0.05078 | 1.000 | − |
|  | R_IFJp | 0.502 | -0.03769 | 1.000 | − |

**Table S3**. The evaluation metrics of the ElasticNet regression on the held-out, out-of-bag (OOB) sub-sample, across 1000 bootstrap iterations, separately for each imaging modality, for the fear acquisition training data. The STACKED row presents the metrics as a result of an additional 500 bootstrap iterations of a second-level multimodal ElasticNet regression model, which received the shortlisted brain areas from all imaging modalities of an earlier 1000-iteration bootstrapped cross-validation in Figure S1. The mean squared error (MSE) and the R² show the goodness of fit and the predictive power of the model on data not seen during the model training.

| **Imaging Modality** | **OOB Mean MSE** | **OOB Median MSE** | **OOB Std. MSE** | **OOB R² Mean** | **OOB Median R²** | **OOB Std. R²** |
| --- | --- | --- | --- | --- | --- | --- |
| VOL | 0.476 | 0.474 | 0.154 | -1.383 | -0.411 | 2.251 |
| SURF | 0.499 | 0.499 | 0.158 | -1.61 | -0.424 | 2.78 |
| THIC | 0.506 | 0.496 | 0.196 | -1.362 | -0.508 | 1.997 |
| INVF | 0.42 | 0.383 | 0.164 | -0.959 | -0.226 | 1.71 |
| ODI | 0.495 | 0.491 | 0.176 | -1.393 | -0.474 | 2.173 |
| RSNE | 0.55 | 0.582 | 0.225 | -1.673 | -0.55 | 2.822 |
| DWINE | 0.505 | 0.516 | 0.168 | -1.477 | -0.533 | 2.33 |
| STACKED | 0.215 | 0.194 | 0.088 | -0.006 | 0.368 | 0.858 |

**Table S4**. The evaluation metrics of the ElasticNet regression on the held-out, out-of-bag (OOB) sub-sample, across 1000 bootstrap iterations, separately for each imaging modality, for the extinction training data. The STACKED row presents the metrics as a result of an additional 500 bootstrap iterations of a second-level multimodal ElasticNet regression model, which received the shortlisted brain areas from all imaging modalities of an earlier 1000-iteration bootstrapped cross-validation in Figure S2. The mean squared error (MSE) and the R² show the goodness of fit and the predictive power of the model on data not seen during the model training.

| **Imaging Modality** | **OOB Mean MSE** | **OOB Median MSE** | **OOB Std. MSE** | **OOB R² Mean** | **OOB Median R²** | **OOB Std. R²** |
| --- | --- | --- | --- | --- | --- | --- |
| VOL | 0.586 | 0.577 | 0.142 | -0.793 | -0.553 | 0.773 |
| SURF | 0.629 | 0.625 | 0.16 | -0.934 | -0.675 | 0.893 |
| THIC | 0.566 | 0.555 | 0.139 | -0.703 | -0.55 | 0.645 |
| INVF | 0.602 | 0.586 | 0.161 | -0.8 | -0.605 | 0.695 |
| ODI | 0.578 | 0.581 | 0.144 | -0.736 | -0.556 | 0.67 |
| RSNE | 0.549 | 0.535 | 0.152 | -0.644 | -0.478 | 0.659 |
| DWINE | 0.566 | 0.562 | 0.142 | -0.696 | -0.543 | 0.621 |
| STACKED | 0.234 | 0.226 | 0.063 | 0.28 | 0.356 | 0.306 |

**Table S5.** Stability of the brain areas across 500 bootstrap iterations for fear acquisition training derived from the second-level stacked model (additional 500 bootstrap iterations), using the pre-filtered region set from the first-level analysis. The **Modality** column represents the imaging modality from which the data have been used to fit the ElasticNet regression. The **Brain Area** column corresponds to brain areas of cortical and, where applicable, subcortical brain areas according to HCPMMP and FreeSurfer's subcortical automated labeling. The ‘L_’ and ‘R_’ prefixes to region codes correspond to the left hemisphere and the right hemisphere. The table with longer anatomical names of the corresponding coded areas is presented at the end of the supplementary materials, Table S7. The **Stability %** shows the percentage of bootstrap iterations where this specific brain area has received a non-zero weight as a result of nested 5-fold cross-validation in predicting a differential (CS+ - CS–) SCR outcome of fear learning. The **Mean Coefficient**, **Sign Consistency**, and **Sign Directionality** show the average beta coefficient of the regularized ElasticNet Regression, consistency of sign, and the directionality (positive or negative) of the beta weight across 5-fold cross-validation in each of 500 bootstrap folds. The rows are sorted by the **Stability %** column in decreasing order for each of the imaging modalities. The list of brain areas is thresholded with at least 50% stability across 500 bootstrap iterations (or being consistently nonzero in at least 250 iterations).

| **Modality** | **Brain Area** | **Stability %** | **Mean Coefficient** | **Sign Consistency** | **Sign Directionality** |
| --- | --- | --- | --- | --- | --- |
| VOL | R_PreS | 0.912 | -0.05595 | 1.000 | − |
|  | R_V3CD | 0.838 | 0.06251 | 0.990 | + |
|  | L_PCV | 0.760 | 0.04615 | 1.000 | + |
|  | R_LO2 | 0.746 | -0.04274 | 0.997 | − |
|  | R_p24pr | 0.730 | 0.03887 | 0.997 | + |
|  | L_MIP | 0.698 | -0.03267 | 0.997 | − |
|  | R_LO1 | 0.692 | 0.04757 | 1.000 | + |
|  | L_TPOJ3 | 0.644 | -0.03023 | 0.997 | − |
|  | R_STV | 0.640 | 0.04106 | 0.978 | + |
|  | L_PreS | 0.624 | -0.02586 | 1.000 | − |
|  | L_MI | 0.596 | -0.03493 | 0.997 | − |
|  | R_43 | 0.590 | -0.02499 | 0.959 | − |
|  | R_8Av | 0.544 | -0.03962 | 0.952 | − |
|  | R_a32pr | 0.532 | 0.02461 | 0.955 | + |
|  | R_MIP | 0.526 | 0.02902 | 1.000 | + |
|  | R_STSva | 0.526 | -0.02962 | 0.996 | − |
|  | R_H | 0.514 | -0.03332 | 1.000 | − |
|  | R_LBelt | 0.510 | 0.03697 | 0.957 | + |
| SURF | R_MIP | 0.844 | 0.03508 | 1.000 | + |
|  | L_PCV | 0.824 | 0.03723 | 1.000 | + |
|  | R_LBelt | 0.594 | 0.02322 | 0.990 | + |
|  | R_STSda | 0.572 | -0.02850 | 0.986 | − |
|  | R_PGi | 0.562 | -0.02466 | 0.961 | − |
|  | L_55b | 0.558 | 0.02542 | 0.867 | + |
| THIC | R_PoI1 | 0.900 | 0.04772 | 1.000 | + |
|  | L_VMV3 | 0.888 | 0.05388 | 0.989 | + |
|  | L_V6A | 0.746 | 0.03171 | 1.000 | + |
|  | L_IFJp | 0.744 | -0.03048 | 0.995 | − |
|  | L_8BL | 0.740 | -0.02887 | 1.000 | − |
|  | L_PHA3 | 0.738 | -0.04611 | 0.978 | − |
|  | L_FOP2 | 0.690 | -0.04378 | 0.971 | − |
|  | L_10d | 0.672 | 0.02620 | 0.997 | + |
|  | L_FEF | 0.670 | 0.03526 | 0.979 | + |
|  | L_MT | 0.606 | 0.02044 | 0.964 | + |
|  | L_IFJa | 0.602 | 0.02825 | 0.980 | + |
|  | L_H | 0.562 | -0.01756 | 0.740 | − |
|  | L_8Av | 0.560 | -0.02557 | 0.964 | − |
|  | L_8Ad | 0.540 | 0.03119 | 0.978 | + |
|  | L_FFC | 0.510 | 0.02309 | 0.984 | + |
| INVF | L_9a | 0.774 | -0.03464 | 0.997 | − |
|  | R_31pv | 0.752 | -0.03098 | 0.997 | − |
|  | L_8BM | 0.714 | 0.03853 | 0.997 | + |
|  | L_A5 | 0.700 | -0.02491 | 1.000 | − |
|  | L_V3A | 0.700 | 0.03438 | 0.983 | + |
|  | L_8Ad | 0.696 | -0.03791 | 0.991 | − |
|  | L_V6 | 0.682 | -0.02858 | 1.000 | − |
|  | R_10pp | 0.668 | -0.02701 | 0.997 | − |
|  | R_8Av | 0.662 | 0.03517 | 1.000 | + |
|  | L_3a | 0.654 | 0.02773 | 0.954 | + |
|  | L_H | 0.652 | -0.03407 | 0.985 | − |
|  | R_LBelt | 0.650 | 0.03071 | 0.957 | + |
|  | L_SFL | 0.638 | 0.02506 | 1.000 | + |
|  | R_LO2 | 0.626 | -0.02647 | 0.990 | − |
|  | L_TPOJ2 | 0.568 | 0.02783 | 0.926 | + |
|  | L_9m | 0.560 | 0.02918 | 1.000 | + |
|  | R_10v | 0.558 | -0.01863 | 1.000 | − |
|  | R_a24pr | 0.554 | 0.02533 | 1.000 | + |
|  | L_47l | 0.546 | 0.04094 | 0.974 | + |
|  | L_FOP1 | 0.542 | -0.03289 | 0.978 | − |
|  | R_AAIC | 0.542 | 0.03195 | 0.804 | + |
|  | L_A1 | 0.502 | -0.02746 | 0.980 | − |
| ODI | L_V3A | 0.918 | -0.03513 | 1.000 | − |
|  | L_A5 | 0.760 | -0.03139 | 0.989 | − |
|  | L_43 | 0.758 | -0.03350 | 0.995 | − |
|  | L_V8 | 0.756 | 0.03324 | 0.997 | + |
|  | R_AIP | 0.722 | 0.02280 | 0.989 | + |
|  | R_31pv | 0.680 | 0.02648 | 0.979 | + |
|  | L_23c | 0.650 | -0.02751 | 0.972 | − |
|  | L_OFC | 0.648 | 0.02251 | 0.991 | + |
|  | R_5m | 0.638 | -0.02138 | 0.991 | − |
|  | R_TPOJ3 | 0.620 | -0.02533 | 0.990 | − |
|  | L_Ig | 0.612 | 0.02675 | 0.882 | + |
|  | L_LIPd | 0.592 | 0.02201 | 0.983 | + |
|  | L_H | 0.590 | -0.02271 | 0.990 | − |
|  | L_caudate | 0.580 | 0.02128 | 0.979 | + |
|  | R_6v | 0.578 | 0.02800 | 0.969 | + |
|  | L_amygdala | 0.544 | 0.01920 | 0.952 | + |
|  | L_24dv | 0.526 | -0.01426 | 0.787 | − |
|  | L_10v | 0.524 | -0.01893 | 0.981 | − |
| RSNE | R_H | 0.604 | -0.02179 | 0.997 | − |
|  | L_OFC | 0.576 | -0.02418 | 0.983 | − |
| DWINE | L_MST | 0.720 | -0.02955 | 1.000 | − |
|  | R_45 | 0.704 | 0.02652 | 0.989 | + |
|  | L_FST | 0.644 | -0.02541 | 1.000 | − |
|  | R_IPS1 | 0.606 | -0.02252 | 0.983 | − |
|  | L_a24pr | 0.590 | 0.02305 | 0.942 | + |
|  | R_47s | 0.590 | -0.02405 | 1.000 | − |
|  | L_VMV2 | 0.578 | 0.02497 | 0.993 | + |
|  | R_V3CD | 0.572 | 0.01979 | 1.000 | + |
|  | R_8Av | 0.570 | -0.01525 | 0.884 | − |
|  | R_STV | 0.560 | 0.02152 | 0.900 | + |
|  | R_9-46d | 0.554 | -0.02439 | 0.993 | − |

**Table S6.** Stability of the brain areas across 500 bootstrap iterations for extinction training derived from the second-level stacked model (additional 500 bootstrap iterations), using the pre-filtered region set from the first-level analysis. The **Modality** column represents the imaging modality from which the data have been used to fit the ElasticNet regression. The **Brain Area** column corresponds to brain areas of cortical and, where applicable, subcortical brain areas according to HCPMMP and FreeSurfer's subcortical automated labeling. The 'L_' and 'R_' prefixes to region codes correspond to the left hemisphere and the right hemisphere. The table with longer anatomical names of the corresponding coded areas is presented at the end of the supplementary materials, Table S7. The **Stability %** shows the percentage of bootstrap iterations where this specific brain area has received a non-zero weight as a result of nested 5-fold cross-validation in predicting a difference of a difference ((Early CS+ - Early CS–) - (Late CS+ - Late CS–)) SCR outcome during extinction learning. The **Mean Coefficient**, **Sign Consistency**, and **Sign Directionality** show the average beta coefficient of the regularized ElasticNet Regression, consistency of sign, and the directionality (positive or negative) of the beta weight across 5-fold cross-validation in each of 500 bootstrap folds. The rows are sorted by the **Stability %** column in decreasing order for each of the imaging modalities. The list of brain areas is thresholded with at least 50% stability across 500 bootstrap iterations (or being consistently nonzero in at least 250 iterations).

| **Modality** | **Brain Area** | **Stability %** | **Mean Coefficient** | **Sign Consistency** | **Sign Directionality** |
| --- | --- | --- | --- | --- | --- |
| VOL | R_STV | 0.740 | 0.07187 | 0.997 | + |
|  | L_24dv | 0.738 | 0.04409 | 1.000 | + |
|  | L_MT | 0.656 | 0.03584 | 1.000 | + |
|  | L_LO3 | 0.638 | 0.04015 | 0.997 | + |
|  | R_V3CD | 0.618 | 0.06166 | 0.997 | + |
|  | R_a9-46v | 0.602 | 0.04129 | 1.000 | + |
|  | L_a24pr | 0.566 | 0.03858 | 1.000 | + |
| SURF | R_7PC | 0.648 | -0.03421 | 0.988 | − |
|  | L_LO3 | 0.622 | 0.02533 | 1.000 | + |
|  | R_44 | 0.580 | 0.03342 | 1.000 | + |
|  | L_V3B | 0.504 | -0.02341 | 0.996 | − |
| THIC | R_PoI2 | 0.830 | 0.04244 | 1.000 | + |
|  | L_5mv | 0.810 | -0.04706 | 1.000 | − |
|  | L_p32 | 0.770 | 0.04639 | 1.000 | + |
|  | L_TE1a | 0.768 | -0.04515 | 1.000 | − |
|  | R_AAIC | 0.754 | 0.03751 | 1.000 | + |
|  | L_PGi | 0.748 | 0.04318 | 1.000 | + |
|  | L_VMV3 | 0.734 | 0.05148 | 0.995 | + |
|  | L_POS1 | 0.664 | -0.04093 | 1.000 | − |
|  | L_s32 | 0.652 | -0.02974 | 0.997 | − |
|  | L_a24pr | 0.648 | 0.03068 | 1.000 | + |
|  | R_10pp | 0.620 | -0.03141 | 1.000 | − |
|  | R_PH | 0.608 | -0.03068 | 0.977 | − |
|  | R_V3CD | 0.576 | 0.03717 | 0.993 | + |
|  | L_3a | 0.574 | -0.03074 | 1.000 | − |
|  | L_1 | 0.548 | 0.03360 | 1.000 | + |
|  | L_SCEF | 0.536 | 0.02637 | 0.996 | + |
|  | R_p24 | 0.532 | -0.03074 | 1.000 | − |
|  | L_6v | 0.520 | -0.03102 | 0.992 | − |
|  | R_OP2-3 | 0.512 | 0.02428 | 1.000 | + |
| INVF | R_13l | 0.876 | 0.05047 | 1.000 | + |
|  | L_TPOJ2 | 0.684 | 0.04854 | 1.000 | + |
|  | L_FOP1 | 0.608 | -0.04062 | 0.997 | − |
|  | R_52 | 0.574 | 0.02626 | 0.986 | + |
|  | L_A1 | 0.554 | -0.03181 | 0.996 | − |
|  | L_8Ad | 0.510 | -0.02417 | 1.000 | − |
|  | L_V3A | 0.506 | 0.02841 | 1.000 | + |
| ODI | L_STGa | 0.850 | -0.04827 | 1.000 | − |
|  | R_8Ad | 0.848 | -0.03591 | 1.000 | − |
|  | L_p32 | 0.814 | -0.03629 | 1.000 | − |
|  | R_9-46d | 0.790 | 0.03308 | 1.000 | + |
|  | R_ProS | 0.750 | -0.03870 | 1.000 | − |
|  | R_6mp | 0.730 | 0.04200 | 1.000 | + |
|  | R_8BL | 0.702 | 0.04054 | 1.000 | + |
|  | R_V4 | 0.690 | -0.03042 | 1.000 | − |
|  | L_RI | 0.688 | 0.03514 | 0.988 | + |
|  | L_V7 | 0.684 | -0.04104 | 0.991 | − |
|  | R_V4t | 0.680 | 0.03074 | 0.997 | + |
|  | L_7PL | 0.674 | -0.03117 | 1.000 | − |
|  | R_9m | 0.650 | 0.03408 | 1.000 | + |
|  | R_IFJa | 0.640 | -0.02594 | 1.000 | − |
|  | R_8C | 0.636 | -0.02520 | 1.000 | − |
|  | L_VMV3 | 0.604 | -0.03339 | 1.000 | − |
|  | R_24dd | 0.600 | 0.02534 | 1.000 | + |
|  | R_VMV3 | 0.582 | -0.02684 | 0.997 | − |
|  | L_47s | 0.548 | 0.02982 | 1.000 | + |
| RSNE | L_accumbens-area | 0.622 | 0.02285 | 1.000 | + |
| DWINE | R_VMV3 | 0.692 | -0.03137 | 1.000 | − |
|  | R_IFJp | 0.686 | -0.03891 | 1.000 | − |
|  | R_PGi | 0.680 | -0.02744 | 0.997 | − |
|  | R_TE2p | 0.648 | -0.02641 | 1.000 | − |
|  | L_13l | 0.634 | -0.02811 | 1.000 | − |
|  | L_VMV3 | 0.568 | -0.02115 | 0.996 | − |
|  | R_VMV1 | 0.524 | -0.01913 | 1.000 | − |
|  | L_p24 | 0.502 | 0.01996 | 0.996 | + |

**Table S7**. HCPMMP cortical regions included in the present analysis, showing the abbreviated region label (Region), full anatomical label (Region Long Name), and corresponding cortical division, following the extended Human Connectome Project multimodal parcellation (HCPex) derived from the original HCPMMP v1.0 atlas (Glasser et al., 2016).

| **Region** | **Region Long Name** | **Cortical Division** |
| --- | --- | --- |
| V1 | Primary_Visual_Cortex | Primary_Visual |
| V2 | Second_Visual_Area | Early_Visual |
| V3 | Third_Visual_Area | Early_Visual |
| V4 | Fourth_Visual_Area | Early_Visual |
| IPS1 | IntraParietal_Sulcus_Area_1 | Dorsal_Stream_Visual |
| V3A | Area_V3A | Dorsal_Stream_Visual |
| V3B | Area_V3B | Dorsal_Stream_Visual |
| V6 | Sixth_Visual_Area | Dorsal_Stream_Visual |
| V6A | Area_V6A | Dorsal_Stream_Visual |
| V7 | Seventh_Visual_Area | Dorsal_Stream_Visual |
| FFC | Fusiform_Face_Complex | Ventral_Stream_Visual |
| PIT | Posterior_InferoTemporal_complex | Ventral_Stream_Visual |
| V8 | Eighth_Visual_Area | Ventral_Stream_Visual |
| VMV1 | VentroMedial_Visual_Area_1 | Ventral_Stream_Visual |
| VMV2 | VentroMedial_Visual_Area_2 | Ventral_Stream_Visual |
| VMV3 | VentroMedial_Visual_Area_3 | Ventral_Stream_Visual |
| VVC | Ventral_Visual_Complex | Ventral_Stream_Visual |
| FST | Area_FST | MT + _Complex |
| LO1 | Area_Lateral_Occipital_1 | MT + _Complex |
| LO2 | Area_Lateral_Occipital_2 | MT + _Complex |
| LO3 | Area_Lateral_Occipital_3 | MT + _Complex |
| MST | Medial_Superior_Temporal_Area | MT + _Complex |
| MT | Middle_Temporal_Area | MT + _Complex |
| PH | Area_PH | MT + _Complex |
| V3CD | Area_V3CD | MT + _Complex |
| V4t | Area_V4t | MT + _Complex |
| 1 | Area_1 | SomaSens_Motor |
| 2 | Area_2 | SomaSens_Motor |
| 3a | Area_3a | SomaSens_Motor |
| 3b | Primary_Sensory_Cortex | SomaSens_Motor |
| 4 | Primary_Motor_Cortex | SomaSens_Motor |
| 23c | Area_23c | ParaCentral_MidCing |
| 24dd | Dorsal_Area_24d | ParaCentral_MidCing |
| 24dv | Ventral_Area_24d | ParaCentral_MidCing |
| 5L | Area_5L | ParaCentral_MidCing |
| 5m | Area_5m | ParaCentral_MidCing |
| 5mv | Area_5m_ventral | ParaCentral_MidCing |
| 6ma | Area_6m_anterior | ParaCentral_MidCing |
| 6mp | Area_6mp | ParaCentral_MidCing |
| SCEF | Supplementary_and_Cingulate_Eye_Field | ParaCentral_MidCing |
| 55b | Area_55b | Premotor |
| 6a | Area_6_anterior | Premotor |
| 6d | Dorsal_area_6 | Premotor |
| 6r | Rostral_Area_6 | Premotor |
| 6v | Ventral_Area_6 | Premotor |
| FEF | Frontal_Eye_Fields | Premotor |
| PEF | Premotor_Eye_Field | Premotor |
| 43 | Area_43 | Posterior_Opercular |
| FOP1 | Frontal_Opercular_Area_1 | Posterior_Opercular |
| OP1 | Area_OP1-SII | Posterior_Opercular |
| OP2-3 | Area_OP2-3-VS | Posterior_Opercular |
| OP4 | Area_OP4-PV | Posterior_Opercular |
| 52 | Area_52 | Early_Auditory |
| A1 | Primary_Auditory_Cortex | Early_Auditory |
| LBelt | Lateral_Belt_Complex | Early_Auditory |
| MBelt | Medial_Belt_Complex | Early_Auditory |
| PBelt | ParaBelt_Complex | Early_Auditory |
| PFcm | Area_PFcm | Early_Auditory |
| RI | RetroInsular_Cortex | Early_Auditory |
| A4 | Auditory_4_Complex | Auditory_Association |
| A5 | Auditory_5_Complex | Auditory_Association |
| STGa | Area_STGa | Auditory_Association |
| STSda | Area_STSd_anterior | Auditory_Association |
| STSdp | Area_STSd_posterior | Auditory_Association |
| STSva | Area_STSv_anterior | Auditory_Association |
| STSvp | Area_STSv_posterior | Auditory_Association |
| TA2 | Area_TA2 | Auditory_Association |
| AAIC | Anterior_Agranular_Insula_Complex | Insula_FrontalOperc |
| AVI | Anterior_Ventral_Insular_Area | Insula_FrontalOperc |
| FOP2 | Frontal_Opercular_Area_2 | Insula_FrontalOperc |
| FOP3 | Frontal_Opercular_Area_3 | Insula_FrontalOperc |
| FOP4 | Frontal_Opercular_Area_4 | Insula_FrontalOperc |
| FOP5 | Area_Frontal_Opercular_5 | Insula_FrontalOperc |
| Ig | Insular_Granular_Complex | Insula_FrontalOperc |
| MI | Middle_Insular_Area | Insula_FrontalOperc |
| PI | Para-Insular_Area | Insula_FrontalOperc |
| Pir | Piriform_Cortex | Insula_FrontalOperc |
| PoI1 | Area_Posterior_Insular_1 | Insula_FrontalOperc |
| PoI2 | Posterior_Insular_Area_2 | Insula_FrontalOperc |
| H | Hippocampus | Medial_Temporal |
| PreS | PreSubiculum | Medial_Temporal |
| EC | Entorhinal_Cortex | Medial_Temporal |
| PeEc | Perirhinal_Ectorhinal_Cortex | Medial_Temporal |
| TF | Area_TF | Medial_Temporal |
| PHA1 | ParaHippocampal_Area_1 | Medial_Temporal |
| PHA2 | ParaHippocampal_Area_2 | Medial_Temporal |
| PHA3 | ParaHippocampal_Area_3 | Medial_Temporal |
| PHT | Area_PHT | Lateral_Temporal |
| TE1a | Area_TE1_anterior | Lateral_Temporal |
| TE1m | Area_TE1_Middle | Lateral_Temporal |
| TE1p | Area_TE1_posterior | Lateral_Temporal |
| TE2a | Area_TE2_anterior | Lateral_Temporal |
| TE2p | Area_TE2_posterior | Lateral_Temporal |
| TGd | Area_TG_dorsal | Lateral_Temporal |
| TGv | Area_TG_Ventral | Lateral_Temporal |
| PSL | PeriSylvian_Language_Area | TPO |
| STV | Superior_Temporal_Visual_Area | TPO |
| TPOJ1 | Area_TemporoParietoOccipital_Junction_1 | TPO |
| TPOJ2 | Area_TemporoParietoOccipital_Junction_2 | TPO |
| TPOJ3 | Area_TemporoParietoOccipital_Junction_3 | TPO |
| 7AL | Lateral_Area_7A | Superior_Parietal |
| 7AM | Medial_Area_7A | Superior_Parietal |
| 7PC | Area_7PC | Superior_Parietal |
| 7Pl | Lateral_Area_7P | Superior_Parietal |
| 7PM | Medial_Area_7P | Superior_Parietal |
| AIP | Anterior_IntraParietal_Area | Superior_Parietal |
| LIPd | Area_Lateral_IntraParietal_dorsal | Superior_Parietal |
| LIPv | Area_Lateral_IntraParietal_ventral | Superior_Parietal |
| MIP | Medial_IntraParietal_Area | Superior_Parietal |
| VIP | Ventral_IntraParietal_Complex | Superior_Parietal |
| IP0 | Area_IntraParietal_0 | Inferior_Parietal |
| IP1 | Area_IntraParietal_1 | Inferior_Parietal |
| IP2 | Area_IntraParietal_2 | Inferior_Parietal |
| PF | Area_PF_Complex | Inferior_Parietal |
| PFm | Area_PFm_Complex | Inferior_Parietal |
| PFop | Area_PF_Opercular | Inferior_Parietal |
| PFt | Area_PFt | Inferior_Parietal |
| PGi | Area_PGi | Inferior_Parietal |
| PGp | Area_PGp | Inferior_Parietal |
| PGs | Area_PGs | Inferior_Parietal |
| 23d | Area_23d | Posterior_Cingulate |
| 31a | Area_31a | Posterior_Cingulate |
| 31pd | Area_31pd | Posterior_Cingulate |
| 31pv | Area_31p_ventral | Posterior_Cingulate |
| 7m | Area_7m | Posterior_Cingulate |
| d23ab | Area_dorsal_23_a + b | Posterior_Cingulate |
| DVT | Dorsal_Transitional_Visual_Area | Posterior_Cingulate |
| PCV | PreCuneus_Visual_Area | Posterior_Cingulate |
| POS1 | Parieto-Occipital_Sulcus_Area_1 | Posterior_Cingulate |
| POS2 | Parieto-Occipital_Sulcus_Area_2 | Posterior_Cingulate |
| ProS | ProStriate_Area | Posterior_Cingulate |
| RSC | RetroSplenial_Complex | Posterior_Cingulate |
| v23ab | Area_ventral_23_a + b | Posterior_Cingulate |
| 10r | Area_10r | AntCing_MedPFC |
| 10v | Area_10v | AntCing_MedPFC |
| 25 | Area_25 | AntCing_MedPFC |
| 33pr | Area_33_prime | AntCing_MedPFC |
| 8BM | Area_8BM | AntCing_MedPFC |
| 9m | Area_9_Middle | AntCing_MedPFC |
| a24 | Area_a24 | AntCing_MedPFC |
| a24pr | Anterior_24_prime | AntCing_MedPFC |
| a32pr | Area_anterior_32_prime | AntCing_MedPFC |
| d32 | Area_dorsal_32 | AntCing_MedPFC |
| p24 | Area_posterior_24 | AntCing_MedPFC |
| p24pr | Area_Posterior_24_prime | AntCing_MedPFC |
| p32 | Area_p32 | AntCing_MedPFC |
| p32pr | Area_p32_prime | AntCing_MedPFC |
| pOFC | Posterior_OFC_Complex | AntCing_MedPFC |
| s32 | Area_s32 | AntCing_MedPFC |
| 10d | Area_10d | OrbPolaFrontal |
| 10 pp | Polar_10p | OrbPolaFrontal |
| 11l | Area_11l | OrbPolaFrontal |
| 13l | Area_13l | OrbPolaFrontal |
| 47m | Area_47m | OrbPolaFrontal |
| 47s | Area_47s | OrbPolaFrontal |
| a10p | Area_anterior_10p | OrbPolaFrontal |
| OFC | Orbital_Frontal_Complex | OrbPolaFrontal |
| p10p | Area_posterior_10p | OrbPolaFrontal |
| 44 | Area_44 | Inferior_Frontal |
| 45 | Area_45 | Inferior_Frontal |
| 47l | Area_47l_(47_lateral) | Inferior_Frontal |
| a47r | Area_anterior_47r | Inferior_Frontal |
| IFJa | Area_IFJa | Inferior_Frontal |
| IFJp | Area_IFJp | Inferior_Frontal |
| IFSa | Area_IFSa | Inferior_Frontal |
| IFSp | Area_IFSp | Inferior_Frontal |
| p47r | Area_posterior_47r | Inferior_Frontal |
| 46 | Area_46 | Dorsolateral_Prefrontal |
| 8Ad | Area_8Ad | Dorsolateral_Prefrontal |
| 8Av | Area_8Av | Dorsolateral_Prefrontal |
| 8BL | Area_8B_Lateral | Dorsolateral_Prefrontal |
| 8C | Area_8C | Dorsolateral_Prefrontal |
| 9-46d | Area_9-46d | Dorsolateral_Prefrontal |
| 9a | Area_9_anterior | Dorsolateral_Prefrontal |
| 9p | Area_9_Posterior | Dorsolateral_Prefrontal |
| a9-46v | Area_anterior_9-46v | Dorsolateral_Prefrontal |
| i6-8 | Inferior_6-8_Transitional_Area | Dorsolateral_Prefrontal |
| p9-46v | Area_posterior_9-46v | Dorsolateral_Prefrontal |
| s6-8 | Superior_6-8_Transitional_Area | Dorsolateral_Prefrontal |
| SFL | Superior_Frontal_Language_Area | Dorsolateral_Prefrontal |
